# Post-inference Methods of Prior Knowledge Incorporation in Gene Regulatory Network Inference

**DOI:** 10.1101/122341

**Authors:** Ajay Nair, Madhu Chetty

**Affiliations:** IITB-Monash Research Academy, Indian Institute of Technology Bombay, Mumbai, India; Chemical Engineering Department, Indian Institute of Technology Bombay, Mumbai, India; Faculty of Information Technology, Monash University, Melbourne, Australia; Faculty of Science and Technology, Federation University, Victoria, Australia

## Abstract

The regulatory interactions in a cell control cellular response to environmental and genetic perturbations. Gene regulatory network (GRN) inference from high-throughput gene expression data helps to identify unknown regulatory interactions in a cell. One of the main challenges in the GRN inference is to identify complex biological interactions from the limited information contained in the gene expression data. Using prior biological knowledge, in addition to the gene expression data, is a common method to overcome this challenge. However, only a few GRN inference methods can inherently incorporate the prior knowledge and these methods are also not among the best-ranked in benchmarking studies.

We propose to incorporate the prior knowledge after the GRN inference so that any inference method can be used. Two algorithms have been developed and tested on the well studied *Escherichia coli*, yeast, and realistic in silico networks. Their accuracy is higher than the best-ranking method in the latest community-wide benchmarking study. Further, one of the algorithms identifies and removes wrong interactions predicted by the inference methods. With half of the available prior knowledge of interactions, around 970 additional correct edges were obtained and 1300 wrong interactions were removed. Moreover, the limitation that only a few GRN inference methods can incorporate the prior knowledge is overcome. Therefore, a post-inference method of incorporating the prior knowledge improves accuracy, removes wrong edges, and overcomes the limitation of GRN inference methods.

## 1 Introduction

Reconstructing the gene regulatory network (GRN) structure from the gene expression data is also known as GRN inference, top-down approach, or reverse engineering of the GRN. Many methods for the GRN inference are available (de Jong, 2002; Markowetz and Spang, 2007). However, the GRN inference is a hard problem due to various challenges, especially due to the limited quantity and quality of experimental data (De Smet and Marchal, 2010; Marbach et al., 2010). A popular method to overcome this limitation is to incorporate prior biological knowledge about the network (expert knowledge) along with the data (Chasman et al., 2016; Ghanbari et al., 2015; Gao and Wang, 2011; Hecker et al., 2009; Cooke et al., 2009; Hartemink et al., 2002). Bayesian network methods are the best known to include the prior knowledge along with the data (Ghanbari et al., 2015; Cantone et al., 2009; Mukherjee and Speed, 2008; Bansal et al., 2007; Gardner and Faith, 2005; Friedman, 2004). In these methods, the prior knowledge is transformed for integration with the scoring function, such as the prior probabilities (Mukherjee and Speed, 2008) or the edge copy number in sampling distribution (Gao and Wang, 2011), ‘before’ the actual inference. These are the best studied methods and we term them as the ‘priors before inference’ methods. There is also a Bayesian network method (Nair et al., 2015) that considers the prior knowledge ‘during’ the inference procedure and not before the inference. This method incorporates the knowledge of the power-law of indegree distribution in GRN network topology, in which only the genes that show a tendency to have a large number of regulators are searched with a larger number of regulators. The number of possible regulators for a gene is not given a priori but is identified ‘during’ the inference from the experimental data. Thus, we term it the ‘prior during inference’ method. In a large-scale assessment of 29 GRN inference techniques (Marbach et al., 2010), a Bayesian network method came in second place. However, the most recent large scale assessment with 35 GRN inference techniques (Marbach et al., 2012) found that the Bayesian network methods scored low when compared to the other methods. Thus, Bayesian network methods are not among the current best-performing GRN inference methods.

A few recent studies reported the inclusion of the prior knowledge with non-Bayesian network methods (Petri et al., 2015; Ellwanger et al., 2014; Chen et al., 2014; Greenfield et al., 2013), in which the prior knowledge is again considered ‘before’ the inference process. Some (Ellwanger et al., 2014; Chen et al., 2014), assume that the GRN structure is available from the prior knowledge and compute only the weight of interactions from the gene expression data. These studies cannot infer any new interactions and thus, cannot be strictly referred to as GRN inference techniques. Petri et al. (2015) uses a supervised machine learning approach to identify new interactions, given the known interactions. Their approach is applicable only when all the regulators are known and at least one target-gene is known for each of these regulators. Greenfield et al. (2013) used a regression method and assumed that all the regulators are known. Assuming that all regulators are known is a hard assumption which prevents discovery of any new regulators. Petralia et al. (2015) reported the integration of different types of experimental data in GENIE3 (Huynh-Thu et al., 2010). GENIE3 is a random forest based GRN inference method which was the best-performer in DREAM5 network inference challenge (Marbach et al., 2012) and DREAM4 in silico network inference sub-challenge (Huynh-Thu et al., 2010). However, only the different types of gene expression data have been integrated and not the prior biological knowledge. Thus, the higher-ranked GRN inference methods are not known to incorporate the prior biological knowledge which can improve their accuracy.

Higher-ranked and other general GRN inference methods are not known to inherently incorporate the prior knowledge into the scoring function and thus, cannot incorporate the prior knowledge along with the experimental data. We hypothesise that this limitation can be overcome if the prior knowledge is incorporated in a post-inference step, after the GRN inference is completed. Since, the prior knowledge is incorporated after the inference, any GRN inference method can be used. Further, it is reported that the methods of reverse engineering the GRN can predict only influence networks (Emmert-Streib et al., 2014; Matos Simoes et al., 2013; Hecker et al., 2009; Gardner and Faith, 2005) because the actual biological interactions cannot be differentiated from the indirect relations between genes. With the post-inference step of prior knowledge inclusion, it could be possible to differentiate the indirect relations from the actual interactions. Since, the prior biological knowledge is incorporated in a post-inference step, we call this as the ‘prior after inference’ method.

This paper has two main contributions: a) two algorithms are developed for the post-inference incorporation of the prior knowledge; and b) a method to identify and remove the indirect interactions using the prior knowledge is reported. The algorithms have been studied using *Escherichia coli* and *Saccharomyces cerevisiae* real networks and also a realistic in silico network, all from the latest benchmarking study, DREAM5 (Marbach et al., 2012). The results are compared to the best-ranked method in the DREAM5.

## 2 Theory and methods

### 2.1 Prior inclusion with any GRN inference method

GRN inference techniques based on the Bayesian network and a few other methods only are known to incorporate the prior biological knowledge along with the gene expression data (Petri et al., 2015; Nair et al., 2015; Greenfield et al., 2013; Gao and Wang, 2011; Cantone et al., 2009; Mukherjee and Speed, 2008; Friedman, 2004; Gardner and Faith, 2005; Bansal et al., 2007). However, these methods were either not considered or did not rank high in the recent DREAM5 benchmarking study (Marbach et al., 2012). Further, their prior knowledge incorporation procedure is specific to the GRN inference technique used such as the prior probabilities (Mukherjee and Speed, 2008), the edge copy number in sampling distribution (Gao and Wang, 2011), or the specific search process (Nair et al., 2015). So, these prior incorporation procedures cannot be readily used with other GRN inference methods. Moreover, the top-ranked methods in DREAM5 are not reported to incorporate the prior knowledge.

In the methods proposed here, the GRN inference is performed using any inference technique and an inferred network is obtained. Then, the post-inference step of incorporating the prior knowledge is performed on the inferred network. Thus, incorporation of the prior knowledge after the inference allows GRN inference using any inference method.

### 2.2 Avoiding inference bias due to prior knowledge

The most common method reported in literature for incorporating the prior knowledge is to include them before the inference (Gao and Wang, 2011; Hecker et al., 2009; Cooke et al., 2009; Cantone et al., 2009; Mukherjee and Speed, 2008; Hartemink et al., 2002). In this case, the inference accuracy is improved by biasing the search with procedures such as formulating a better initial set of hypothesis based on the prior knowledge (Mukherjee and Speed, 2008) or restricting the search space towards solutions that include the prior knowledge (Gao and Wang, 2011; Hecker et al., 2009). A problem with this method is that if an interaction in the prior knowledge is not present in the experimental data (due to the experimental conditions being different or due to the lack of transcript correlation between regulator and its target-gene), the use of the prior knowledge will still bias the search. This bias occurs because, inferring one interaction can influence the inference of subsequent interactions. The subsequent interactions can be missed or added wrongly in heuristic search methods such as greedy search or in the optimal methods that use decomposable scoring. Thus, incorporating the prior knowledge not supported by data may lead to regulatory interactions supported by the data to be missed. This is not desirable because the predicted network can be unnecessarily inaccurate due to the prior knowledge.

In the prior after inference method proposed here, at first, the inference is performed on the data. Thus, a network that is a true representation of the data is first obtained. Subsequently, in the post-inference step, the prior knowledge is incorporated in the obtained network which will not bias the inference process.

### 2.3 Identification and removal of wrong edges

It is reported that the GRN inference from the gene expression data infers only an influence network which contains transcription regulatory interactions along with other regulatory and non-regulatory interactions (Emmert-Streib et al., 2014; Matos Simoes et al., 2013; Hecker et al., 2009; Gardner and Faith, 2005). However, while quantifying the accuracy of the network inference, only the transcriptional regulatory interactions are considered as correct edges or true positives. All other interactions are considered to be wrong edges or false positives. It has been observed that even high-ranking methods in benchmark studies (Marbach et al., 2012) show very low precision which means that they predict a significantly high number of wrong edges. We identify the source of these wrong edges and develop a method to reduce them.

#### 2.3.1 Sources of wrong edges

A GRN shows the interactions between regulators and their target-genes. The complex bio-physical process by which a regulatory transcription factor protein controls the gene expression of a target-gene is represented as an edge between the regulator and the target-gene in a GRN. The gene expression data measures concentration of mRNA transcripts in cells. However, inferring the GRN from the gene expression data can detect only statistical dependencies between the mRNA transcripts in the gene expression data.

The dependencies between the mRNA transcripts can be due to interactions at transcription, post-transcription, protein, and metabolite (Emmert-Streib et al., 2014; Friedman, 2004; Gardner and Faith, 2005) which are not measured separately. Thus, the non-transcriptional interactions can be inferred due to the information limitation in the data. It is reported that the GRN inference methods based on the gene expression data can infer only an influence network (Emmert-Streib et al., 2014; Matos Simoes et al., 2013; Hecker et al., 2009; Gardner and Faith, 2005). The influence networks can contain any interaction that influences the mRNA expression levels such as the direct and indirect transcriptional regulatory interactions (Friedman, 2004); or other non-transcriptional regulatory and non-regulatory interactions such as the protein-protein interactions and metabolite interactions (Gardner and Faith, 2005). Thus, the inferred network may contain direct and indirect transcriptional regulatory interactions and other influence interactions.

Another challenge in using the gene expression data is that the regulator and its target-gene mRNA transcript concentration are not always correlated (Chua et al., 2004). From our understanding of the gene regulation, the regulatory proteins control the expression of its target-gene mRNA. If this is true then the lack of correlation could be because the regulatory protein activity is not correlated to its mRNA concentration and/or the regulator mRNA concentration does not vary. Both of these conditions have been observed. It has been reported that the activity of the regulators do not necessarily correlate with their transcript abundance (Hecker et al., 2009; Gardner and Faith, 2005; Herrgard et al., 2003). This could be because the activity of some regulatory proteins is controlled by their phosphorylation states. Phosphoproteome studies in yeast (Pomraning et al., 2016) have shown that around 18% of the regulators have at least one phosphorylation domain by which they are activated or deactivated. Moreover, it is also reported that most regulators are themselves not transcriptionally regulated and have a low constitutive mRNA expression (Herrgard et al., 2003). Thus, even if a regulator regulates a target-gene, it may not be detected at the gene expression level of their transcripts. This lack of correlation between some of the regulators and their target-gene mRNAs will result in some correct interactions being undetected.

If different target-genes are regulated by the same regulatory proteins, then the target-gene transcripts can be correlated. A condition in which the correlation between the regulator and the target-gene transcripts are absent but correlation between the target-gene transcripts are present, can lead to the inference of interactions between the target-genes. These inferred interactions between the target-genes do not represent real biological interactions but indirect transcriptional interactions due to the common regulator. It is reported that the indirect interactions can also produce statistical dependencies (Friedman, 2004; Gardner and Faith, 2005). Thus, the indirect transcriptional interactions can also be present in an inferred network.

Other factors such as noisy, limited, and incomplete data or errors in experiments can cause spurious interactions to be inferred (Emmert-Streib et al., 2014; Gardner and Faith, 2005). Thus, due to the limitations in technology or experimental errors, the inference may not capture the actual biological interaction or may predict spurious interactions.

Therefore, the three known reasons that cause the wrong edges or the false positives in an inferred network are the influence interactions, the indirect interactions and the spurious interactions. The influence interactions are the protein or metabolite level interactions (which are not transcription regulatory) that influence the gene expression. The indirect interactions are the interactions inferred due to the statistical dependencies shown by the indirect transcriptional regulatory interactions. The spurious interactions are the interactions inferred due to noise, errors, or such limitations in data or experiments.

#### 2.3.2 Analysis of wrong edges

Previous studies (Yu et al., 2004; Marbach et al., 2010) have analysed the wrong edges or the false positives predicted by different types of GRN inference methods. It is reported that these false positives have a certain pattern. If the inferred network is visualised as a graph, then the false positives either originate from indirectly connected genes (known as relatives) or far away genes (known as strangers). Since the relatives are defined with respect to the GRN, the false positives from the relatives originate from the indirect interactions that are defined in section 2.3.1. Yu et al. (2004) showed that in Bayesian network methods the false positives from relatives are significantly large and for larger data, the false positives from the relatives are greater than the strangers. They further classified the false positives from relatives into grandparent, uncle, sibling, or child. Marbach et al. (2010) analysed results from 29 GRN inference methods and reported that these methods suffer from systematic prediction errors in the network motifs that involve relatives. For example, fan-out error showed the false positives caused by siblings and cascade error showed the false positives caused by grandparents.

Yu et al. (2004) classified the sources of false positives as grandparent, sibling uncle, child, and stranger. In their classification, a three-node distance in direction of ancestor was considered such as the uncle, while in the direction of descendants only a one-node distance, the child, was considered. Our analysis show that the false positives also occur from three-node distance descendant, the grandchildren. Further, they considered only four types of relatives and all other relatives were grouped with the strangers. This approach is not intuitive and no justification was provided. Therefore, we expanded the classification to include the same node distances in the ancestor and descendant directions.

The previous study (Yu et al., 2004) considered three-node distance relatives. We consider up to four-node distance relatives in ancestor and descendant directions. The relatives beyond the four-node distance are not considered because it is not possible to detect them when the prior knowledge is limited. Moreover, the number of false positives from them is also negligible. The details of the relatives and strangers up to four-node distances are given in figure 1. In the figure, ‘TG’ is the target-gene whose relatives are illustrated. For example, ‘Prnt’ is the regulator or the parent of the TG and ‘Strg’ represents the possible strangers within the four-node distance. Other relatives are also defined with respect to the TG. The number on the edges, in the figure, represents the node distance from the TG. The details of the different relatives of the TG, their node distance, and their abbreviations used are as follows:

- Distance 1: child (‘Chd’)
- Distance 2: sibling (‘Sib’), grandparent (‘Gp’), grandchild (‘Gc’)
- Distance 3: aunt (‘Aun’), niece (‘Nie’), great-grandparent (‘GGp’), great-grandchild (‘GGc’)
- Distance 4: cousin (‘Cus’), child-of-niece or Niece-child (‘NCd’), grandparent’s sibling (‘GpS’), great-great-grandparent (‘GGGp’), great-great-grandchild (‘GGGc’)

**Figure 1:**
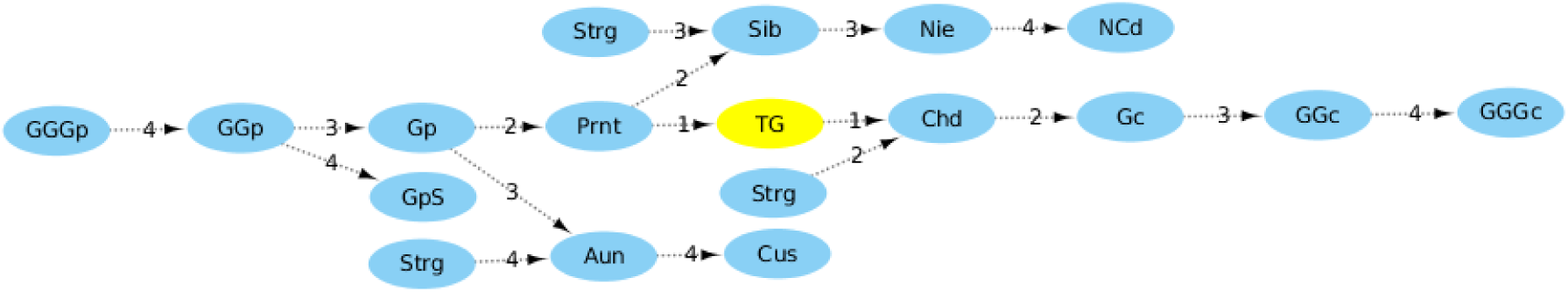
The different relatives of the TG, up to four node distance, are represented in a directed graph. The names of the nodes are defined in the text. The numbers on the arrows represent the node distance from the TG.

The false positives from the relatives will be termed as relative-FPs (relative-false-positives). Similarly, the false positives from each relative will be denoted with an ‘FP’ suffixed to the relation name such as sibling-FP, grandparent-FP, aunt-FP and so on. The main challenge is that a relative-FP can fall into more than one relationship category. For example, let TG1→TG2 be an inferred edge that is found to be a false positive, based on the prior knowledge. Further, it is known from the prior knowledge that TF1→TG1, TF1→TG2, TF2→TF1, and TF2→TG1 interactions exist. Then, TG1 can be a sibling-FP or an aunt-FP to TG2. To overcome this multiple category problem, we use a ranking for relatives based on the node distance because influence is larger from shorter distances. Further, the ancestors are higher ranked than descendants because the ancestors’ influence will be causal, while the descendants show a non-causal correlation. Thus, the following preference order is followed in classifying the false positives: sibling (highest rank), grandparent, aunt, niece, great-grandparent, cousin, niece-child, grandparent-sibling, great-great-grandparent, child, grandchild, great-grandchild, and great-great-grandchild (lowest rank). Therefore, according to the preference order, if a node is both a sibling-FP and an aunt-FP, it will be classified as a sibling-FP.

The previous studies (Yu et al., 2004; Marbach et al., 2010) have reported that different GRN inference methods predict similar types of false positives. Many false positives originate from the relatives (relative-FPs) and rest from the strangers. It is shown here that the relative-FPs are caused mainly due to the indirect interactions. Further, we propose to classify the relative-FPs into 13 types of false positives.

#### 2.3.3 Removal of indirect interactions

We hypothesise that, if the knowledge of actual interactions (prior knowledge) is available, then it would be possible to identify and remove the relative-FPs or the indirect interactions. For example, let Sib→TG be an edge in the inferred network. In the prior knowledge, let: a) Prnt→TG, b) Prnt→Sib edges exist, and c) Sib not be a regulator (no DNA binding domain). Then, it can be deduced that the Sib→TG edge is an indirect interaction or a relative-FP. Thus, by incorporating the prior knowledge in a post-inference step, a) the actual interactions Prnt→TG and Prnt→Sib can be included, and b) the sibling-FP (Sib→TG) indirect interaction in the inferred network can be identified and removed.

The prior knowledge of transcriptional interactions (edge priors, non-edge priors, and known regulators) may detect only the indirect interactions as shown by the previous example. The influence edges due to protein-protein interactions or metabolites may not be detected because these prior information are not considered. However, some influence interactions may get removed. For example, let G1→G2 be an influence edge inferred due to G1 and G2 protein interactions. If the prior knowledge states that G1 cannot be a regulator, then this influence edge will be removed. Thus, using the prior knowledge, the indirect interactions and possibly other wrong edges can be removed.

#### 2.3.4 Conditions for identifying indirect interactions

In the previous section, the condition for identifying an indirect interaction from a sibling was described. Here, the conditions required for identifying the indirect interactions from the different relatives are developed. Figure 2 shows the inferred wrong interaction PInf→TG (edge label ‘a’) and all the possible relatives from which the correct indirect interaction could have originated. PInf is the parent-inferred or the inferred regulator of the TG that is not its correct parent or the regulator.

**Figure 2:**
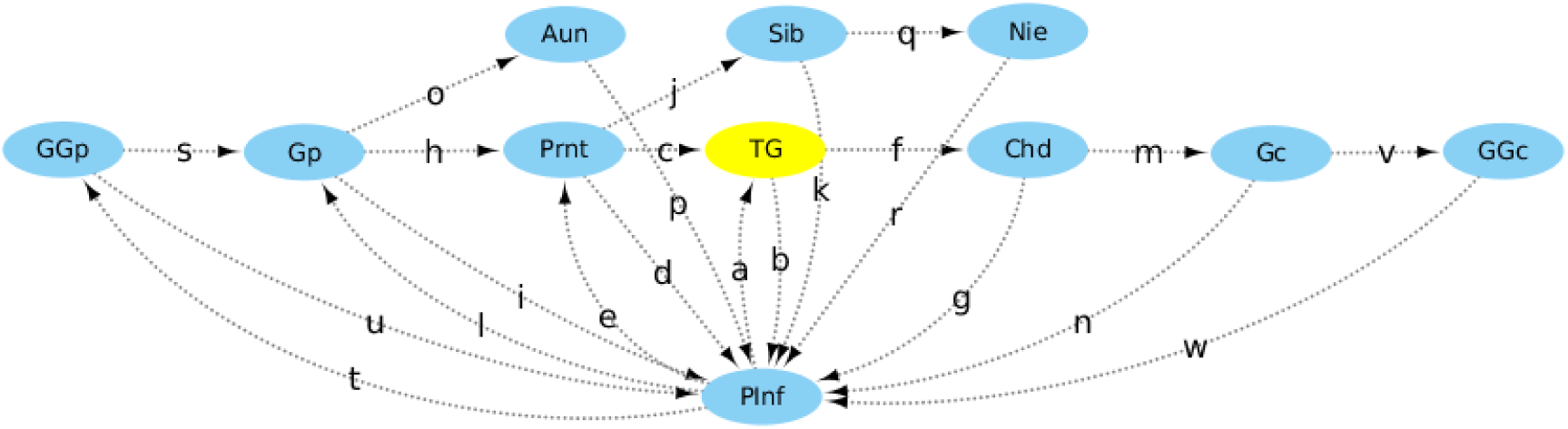
The inferred wrong interaction PInf→TG and all its possible indirect interactions. PInf is wrongly the parent-inferred or the inferred regulator for the node TG. The network shows all possible relatives that the PInf can be, for the node TG. Relatives up to four-node distances only are considered.

The conditions for identifying the relative-FPs are given in Table 1. As an example, consider the identification of niece-FP. In the niece-FP condition, the PInf→TG edge has been wrongly inferred and actually the PInf is a niece to the TG. This mistake can be identified if the following edges and non-edges are present in the prior knowledge: edges *c*,*j*,*k* are present as edge priors and edges *a*,*b*,*d*,*e*, (*fg*), (*hi*) are known to be absent (non-edge priors). When edges *c, j, k* are present, a parent, a sibling, and a niece are present, respectively, for the TG and the PInf is its niece. The absence of edges *a* and b signifies that the nodes TG and PInf do not interact directly. The absence of edges d and e implies that the PInf is not a sibling or a grandparent, respectively, of the TG. Further, the absence of (*fg*) (where either edges f and *g*, or both are absent) implies that the PInf is not a descendant of the TG. The absence of (*hi*) implies that the PInf is not an aunt of the TG. Thus, the closest relation of the PInf to the TG is a niece. In Table 1, the nomenclature a′ means absence of edge *a* and (*fg*)′ means either of edges *f* and *g*, or both of them are absent. Thus, the conditions given in Table 1 identify the indirect interactions up to four-node distance.

**Table 1:** Conditions for classifying the false positives from relatives (relative-FPs) for up to a four-node distance. The edge and non-edge labels are from figure 2

| Distance | Relation | Edges | Non-edges |
| --- | --- | --- | --- |
| - | True positive | a | - |
|  | False positive (FP) | - | a' |
| 1-node | Child-FP | b | a' |
| 2-node | Sibling-FP | c, d | a', b' |
|  | Grandparent-FP | c, e | a', b', d' |
|  | Grandchild-FP | f, g | a', b', (cd)', (ce)' |
| 3-node | Aunt-FP | c, h, i | a', b', d', e', (fg)' |
|  | Niece-FP | c, j, k | a', b', d', e', (fg)', (hi)' |
|  | Great-grandparent-FP | c, h, l | a', b', d', e', (fg)', i', (jk)' |
|  | Great-grandchild-FP | f, m, n | a', b', (cd)', (ce)', (cjk)', (chi)', (chl)', g' |
| 4-node | Cousin-FP | c, h, o, p | a', b', d', e', (fg)', i', (jk)', l', (fmn)' |
|  | Niece-child-FP | c, j, q, r | a', b', d', e', (fg)', (fmn)', (hi)', (hl)', (hop)', k' |
|  | Grandparent-sibling-FP | c, h, s, u | a', b', d', e', (fg)', i', (jk)', l', (fmn)', (jqr)', (op)' |
|  | Great-great-grandparent-FP | c, h, s, t | a', b', d', e', (fg)', i', (jk)', l', (fmn)', (jqr)', (op)', u' |
|  | Great-great-grandchild-FP | f, m, v, w | a', b', (cd)', (ce)', (cjk)', (chi)', (chl)', g', (cjqr)', (chop)', (chst)', (chsu)', n' |

### 2.4 Advantages of prior after inference

Incorporating the prior knowledge in a post-inference step has the following advantages: a) it improves the accuracy of inference due to the incorporation of the prior knowledge (Steele et al., 2009; Gao and Wang, 2011; Isci et al., 2014); b) it identifies and removes the wrong edges due to the indirect interactions; c) it helps to incorporate the prior knowledge with any GRN inference technique and overcomes the limit in choice of inference methods; d) it avoids the inference process bias towards inaccurate results due to the prior knowledge; and e) it shortlists the edge priors not supported by the data for further study.

Section 2.8 discusses the different reasons that can cause an edge prior to be not supported by the gene expression data. These edge priors are interesting candidates for further study in wet-lab. Although, any method of prior incorporation can check if the priors are supported by the data, this step is more intuitive in the prior after inference method because the design philosophy of the method considers the possibility of the lack of transcript correlation between a regulator and its target-gene, noise, errors, and so on.

### 2.5 Algorithm development

The steps required to obtain the full advantages of a prior after inference method are: a) incorporation of the prior knowledge of interactions, b) incorporation of the knowledge of regulators, c) identification and removal of the wrong edges, and d) shortlisting of the prior interactions not supported by the data. Of these, the most basic step and the procedure that is similar to the existing methods of prior knowledge incorporation, is to perform only the first step of incorporating the prior knowledge of interactions. Thus, two prior after inference (post-inference prior) algorithms are proposed. The first, termed the Basic post-inference algorithm, performs only the incorporation of the prior knowledge of interactions. The second, termed the Advanced post-inference algorithm, performs all the four steps.

### 2.6 Basic post-inference algorithm

The Basic post-inference algorithm or the Basic algorithm incorporates the prior knowledge of interactions into the result (the inferred network) obtained from a normal GRN inference process without the prior knowledge. Since this is the most basic form of prior incorporation in a post-inference method, it is termed as the Basic post-inference algorithm. The inputs required for the algorithm are the following.

1. The inferred network which is obtained from any GRN inference method. The inferred network, termed the *infMat*, is an *N* × *N* matrix that represents the pairwise relation between all the *N* number of genes in the network. For any two genes *i* and *j*, the pairwise relation is defined as,

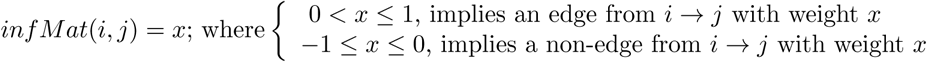

2. The prior knowledge on interaction — the edge and the non-edge priors. These are given as two separate *N* × *N* matrices, *edgePrior* and *nEdgePrior*, respectively. For any two genes *i* and *j*, these matrices are defined as,

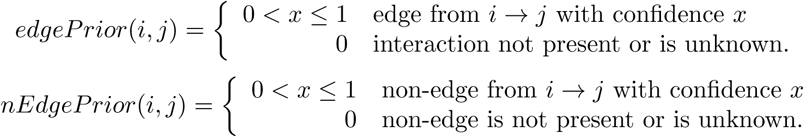

It should be noted that the *edgePrior* and the *nEdgeP rior* are complimentary such that for any two genes *i* and *j*, both *edgePrior*(*i, j*) and *nEdgePrior*(*i, j*) cannot be non-zero simultaneously. Further, *edgePrior*(*i, j*) = 1, implies a full confidence prior edge between genes *i* and *j*, Similarly *nEdgePrior*(*i, j*) = 1, implies a full confidence non-edge prior.

In the Basic algorithm, the GRN inference is performed with any inference method of user’s choice and an inferred network is obtained. The *infMat* is created from the inferred network by simple operations such as scaling to make the weight of interactions between genes to range from −1 to 1. The positive weights represent confidence in the edges between genes, with 1 being the highest confidence. The zero and negative weights indicate the confidence in non-edges, with −1 representing the highest confidence. If weights for the non-edges are not available in the inferred network, then the range of weights in the *infMat* will be between 0 and 1, with 0 representing the absence of edges. For cases in which only the edges and the absence of edges are known and the weight of interactions are not available, the *infMat* will be a binary matrix with 1 and 0 representing an edge and a non-edge, respectively.

Once the *infMat* is obtained, the pseudo-code for rest of the procedure is given in Algorithm 1. In the algorithm, the prior knowledge of edges and non-edges are incorporated for each interacting gene pairs. The edge prior weights in *edgePrior* are added and the non-edge prior weights in *nEdgePrior* are subtracted from the *infMat* weights. The *edgePrior* and the *nEdgeP rior* weights are multiplied by a factor 2 to ensure that an incorrect prediction in the *infMat* is also corrected. For example, assume *infMat*(*i, j*) = −1, which implies a non-edge between genes *i* and *j*. However, if an edge actually exists with *edgePrior*(*i, j*) = 1, then *infMat*(*i, j*) + 2 * *edgePrior*(*i, j*) will result in the edge between *i* and *j* being incorporated in the final network. The final post-inference network that incorporates the prior knowledge into the *infMat* is an *N* × *N* matrix *BPI* (acronym for Basic post-inference); which represents the pairwise relation between all genes after the GRN inference and post-inference prior incorporation. For any two genes *i* and *j*, the pairwise relation in *BPI* is defined as,

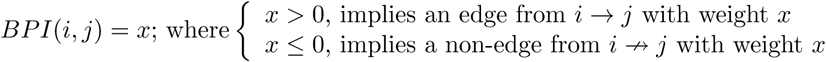

From Algorithm 1, it can be seen that all the possible pairwise interactions are considered for the post-inference process. Since *N* genes are present, the complexity of this algorithm is thus, *O*(*N^2^*).

#### Algorithm 1 Basic post-inference algorithm

Inputs: Inferred network (*infMat*), edge priors (*edgePrior*) and non-edge priors (*nEdgePrior*). Assuming *N* genes are present, these inputs are *N* × *N* matrices.

~~~
                                                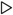**Integrate the prior knowledge with the inferred network**
**for** i=1 to N; **do**
           **for** j=1 to N; **do**
                           *BPI*(*i, j*) = *infMat*(*i, j*) + 2 * *edgePrior*(*i, j*) − 2 * *nEdgeP rior*(*i, j*)
           **end for**
         **end for**
~~~

### 2.7 Advanced post-inference algorithm

The Advanced post-inference algorithm or the Advanced algorithm, incorporates the prior knowledge of interactions, includes the prior knowledge of regulators, identifies and removes the wrong edges, and shortlists the prior interactions not supported by the data. The inputs required for the algorithm are the following.

1. The inferred network, *infMat*, obtained from any GRN inference method. The details of *infMat* are given in section 2.6.
2. The prior knowledge on interaction given by *edgePrior* and *nEdgeP rior*. These matrices are also discussed in section 2.6.
3. The knowledge of the transcription factor and the non-transcription factor genes (for example the genes that lack DNA binding domains cannot be transcription factors), given by a 1 × N matrix—*tf*Possible. For gene *i*, this matrix is defined as,

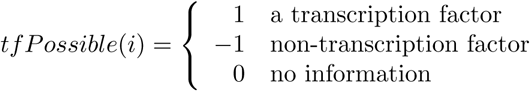
4. The knowledge that all GRN inference methods based on the gene expression data predict indirect interactions that can be identified and removed. These indirect interactions are classified as wrong edges by the performance metrics. It was shown in section 2.3 that the knowledge of actual interactions can be used to identify and remove the wrong edges caused by the indirect interactions.
5. The gene expression data that was used to infer the *infMat* in an *n* × *N* matrix, in which *n* is the number of experiment data samples. This data is used to check the support for the edge priors.

In the Advanced algorithm, similar to the Basic algorithm, the *infMat* is obtained from any GRN inference method. The other user inputs required are *edgePrior, nEdgePrior, tfPossible*, and gene expression data. Once these inputs are obtained, the rest of the procedure is given as a pseudo-code in Algorithm 2. The main steps of the algorithm are: a) find and report the edge priors without data support; b) identify and remove the wrong edges (implemented as two separate steps of ancestor-FP and descendant-FP removal); c) incorporate the prior knowledge of regulators; and d) integrate all the prior knowledge and compute the final network.

In the procedure to find the edge priors without data support, each edge prior is individually checked. Since each edge prior corresponds to a transcription regulatory interaction between two genes, a statistical dependency is expected in the gene expression data between these two genes. The statistical dependency can be measured with tools such as mutual information, the details of which are given in section 2.8. The edge priors that do not show statistical dependency in the gene expression data are identified and reported. The algorithm still includes the non-data support edge priors because the absence of data support can be due to the lack of regulator and target-gene transcript correlation, noise, or error. One of the main focus of the Advanced algorithm is to remove the false positives caused due to these reasons.

The procedure to identify and remove wrong edges is based on the theory developed in section 2.3. The wrong edges from the relatives are termed the indirect interactions (or relative-FPs) which can be identified and removed using the prior knowledge. The conditions to identify the indirect interactions were given in Table 1. In the table, a large number of non-edge priors were required to identify a relative-FP. Non-edge priors are not easily available and though there are methods to obtain them (Nair et al., 2014), it needs organism specific knowledge; which is not feasible for a general method developed here. Further, the quantity of the edge priors available is also generally low. Due to these factors, the Advanced algorithm proposes a pragmatic approach to classify the relative-FPs. Here, the essential edge priors and any available non-edge priors are only considered. Further, the ancestor-FPs are checked first and then the descendant-FPs because only the ancestors have a causal relation. Moreover, because the quantity of priors is limited, the relative-FP classification is performed only for the inferred edges related to the available priors; and not necessarily for all the inferred edges.

The conditions that incorporate these approaches to classify the relative-FPs are given in Table 2. Due to the simplification of the conditions in Table 2 when compared to Table 1, confidence weights of some correct inferred edges that are not in the edge priors could be reduced and some low confidence edges could also be removed, in the post-inference stage. This will be the cost of trade-off for implementing the algorithm when the prior knowledge is limited. However, because only the low confidence inferred edges are expected to be affected, the overall performance of the Advanced algorithm is expected to be better than the Basic algorithm.

**Table 2:** Conditions for classifying the false positive relatives (relative-FPs) in the Advanced algorithm. The relatives up to four-node distances are considered.

| Distance | Relation | Edges | Non-edges |
| --- | --- | --- | --- |
| - | True positive | a | - |
|  | False positive (FP) | - | a' |
| Ancestors |  |  |  |
| 2-node | Sibling-FP | c, d/$\hat{d}$ | a*, b* |
|  | Grandparent-FP | c, e | a*, b*, d* |
| 3-node | Aunt-FP | c, h, i | a*, b*, d*, e* |
| | Niece-FP | c, j, k/$\hat{k}$ | a*, b*, d*, e*, (hi)* |
|  | Great-grandparent-FP | c, h, l | a*, b*, d*, e*, i*, (jk)* |
| 4-node | Cousin-FP | c, h, o, p | a*, b*, d*, e*, i*, (jk)*, l* |
|  | Niece-child-FP | c, j, q, r | a*, b*, d*, e*, (hi)*, (hl)*, (hop)*, k* |
|  | Grandparent-sibling-FP | c, h, s, u | a*, b*, d*, e*, i*, (jk)*, l*, (jqr)*, (op)* |
|  | Great-great-grandparent-FP | c, h, s, t | a*, b*, d*, e*, i*, (jk)*, l*, (jqr)*, (op)*, u* |
| Descendants |  |  |  |
| 1-node | Child-FP | b | a* |
| 2-node | Grandchild-FP | f, g | a*, b* |
| 3-node | Great-grandchild-FP | f, m, n | a*, b*, g* |
| 4-node | Great-great-grandchild-FP | f, m, v, w | a*, b*, g*, n* |

The relative node names, edge labels, and non-edge labels in Table 2 are based on figure 2. In the nomenclature followed in table: a — represents the presence of an edge prior; a’ — represents the presence of a non-edge prior;a* — represents the absence of an edge prior; and â —represents the presence of an inferred edge along with other relevant prior knowledge such as a transcription factor. For example, in the case of niece-FP relation, *PInf* is a niece to the *TG* and the *PInf* → *TG* edge is wrongly inferred. This wrong interaction can be identified if the following conditions are true. a) Edges *c* and *j* are present as edge priors. b) Either edge *k* is present as an edge prior or the three conditions — *k* is not a non-edge prior, *Sib* node is a transcription factor, and *k* is an inferred edge — are true. These three conditions are represented by 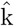 in the table. c) Edges *a, b, d, e*, (*hi*) are absent in the prior knowledge.

#### Algorithm 2 Advanced post-inference algorithm

~~~
1: Inputs: Inferred network (*infMat*), edge priors (*edgePrior*), non-edge priors (*nEdgeP rior*), gene expression data, and regulatory genes *(tfPossible)*.
2: **procedure** Find AND REPORT EDGE PRIORS WITHOUT DATA SUPPORT
3:   Edge priors not supported by data are found using statistical methods such as mutual information.
4: **end procedure**
5: Initialise these *N×N* matrices to zero: *ppEdge,ppNEdge,ppEdgeTF,ppNEdgeTF*
6: **procedure** IDENTIFY AND REMOVE WRONG EDGES (ANCESTOR-FPs)
7:   **for each** *Prnt* → *TG* in *edgePrior* **do**
8:       Update *Prnt → TG* edge in *ppEdge(Prnt, TG)* and *TG 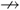 Prnt* non-edge in *ppNEdge(TG, Prnt)*
9:       **for** each *PInf* of the *TG* in *infMat* **do**
10:          If *PInf* → *TG* is a prior edge, update *ppEdge,ppNEdge* appropriately. Skip rest of the steps, and go to next iteration of the current for loop
11:          If *PInf* → *TG* is a non-edge, update *ppNEdge* appropriately.
12:          If *PInf* is a sibling-FP, update *ppEdge,ppNEdge* appropriately.
13:          Else, if *PInf* is a grandparent-FP, update *ppEdge,ppNEdge* appropriately.
14:          Else, if *PInf* is an aunt-FP, update *ppEdge,ppNEdge* appropriately.
15:          Else, if *PInf* is a niece-FP, update *ppEdge,ppNEdge* appropriately.
16:          Else, if *PInf* is a great-grandparent-FP, update *ppEdge,ppNEdge* appropriately.
17:          Else, if *PInf* is a cousin-FP, update *ppEdge,ppNEdge* appropriately.
18:          Else, if *PInf* is a niece-child-FP, update *ppEdge,ppNEdge* appropriately.
19:          Else, if *PInf* is a grandparent-sibling-FP, update *ppEdge,ppNEdge* appropriately.
20:          Else, if *PInf* is a great-great-grandparent-FP, update *ppEdge,ppNEdge* appropriately.
21:        **end for**
22:      **end for**
23:**end procedure**
24:**procedure** Identify and remove wrong edges (descendant-FPs)
25:   **for** each *TG* → *Child* in *edgePrior* **do**
26:       Update *TG* → *Child* edge in *ppEdge(TG, Child)* and *Child → TG* non-edge in *ppNEdge(Child, TG*)
27:       for each *PInf* of the *TG* **do**
28:           If *PInf* → *TG* is a prior edge, then update weights in *ppEdge,ppNEdge*, skip rest of the steps, and go to next iteration of the current for loop
29:           If *PInf → TG* is a non-edge, update *ppNEdge*.
30:           If *PInf* is a child-FP, update *ppEdge,ppNEdge* appropriately.
31:           Else, if *PInf* is a grandchild-FP, update *ppEdge,ppNEdge* appropriately.
32:           Else, if *PInf* is a great-grandchild-FP, update *ppEdge,ppNEdge* appropriately.
33:           Else, if *PInf* is a great-great-grandchild-FP, update *ppEdge,ppNEdge* appropriately.
34:         **end for**
35:      **end for**
36:**end procedure**
37:**procedure** Incorporate the knowledge of regulators
38:   Weights of inferred edges of relevant regulators are updated in *ppEdgeTF*.
39:   Weights of non-edges from non-TFs are updated in *ppNEdgeTF*.
40:**end procedure**
41:**procedure** INTEGRATE PRIOR KNOWLEDGE AND COMPUTE THE FINAL NETWORK
42:   *w* = 2*maximum_value_in(*ppEdge,ppEdgeTF,ppNEdge*).
43:   *API = infMat − ppNEdge* + *ppEdge* + *ppEdgeTF − w* * *ppNEdgeTF* + *w* * *edgePrior − w* * *nEdgePrior*
44:**end procedure**
~~~

In the identify-and-remove-wrong-edges procedure, the weights of known and expected edges are in the *N*×*N* pairwise edge confidence matrix, *ppEdge*. Similarly, the weights of known and expected non-edges are in the *N*×*N* pairwise non-edge confidence matrix, *ppNEdge*. For example, if a relative-FP is identified, the required edge interactions are incorporated by increasing the interaction weights between the appropriate genes in *ppEdge*. Similarly, the required non-edge interactions are incorporated by increasing the appropriate weights in *ppNEdge*. Therefore, after the identify-and-remove-wrong-edges procedure, the *ppEdge* and the *ppNEdge* matrices will have the cumulative confidence of the expected edge and non-edge interactions, respectively.

The next procedure is to incorporate the knowledge of regulators. This procedure consists of two steps: a) increasing the weights of inferred edges from the known transcription factors; and b) removing the inferred wrong edges from the non-transcription factors. The prior knowledge of transcription factors and non-transcription factors are in the variable *tfPossible*. If the *infMat* has inferred edges from the known transcription factors, these are more likely to be true edges because the GRN inference methods are designed to identify such interactions. Moreover, because the regulator and its target-gene transcript correlation can be low (discussed in section 2.3), these inferred edges may have lower confidence weights. Due to these two reasons, the confidence weights of the inferred edges from the known transcription factors in *infMat* are increased. An additional constrain, that these known transcription factors should also regulate a gene among the considered genes, is kept to prevent false interactions. Thus, in the *infMat*, for each inferred edge from a known transcription factor that also has a known target-gene, the corresponding pairwise interaction weight is increased in the *N* × *N* matrix, *ppEdgeTF*. It was shown in section 2.3 that wrong edges can be inferred from the non-transcription factors. For a non-transcription factor in the *tfPossible*, any edge from it to another gene will be a wrong edge and should be removed. The information of edges going out of each non-transcription factor is updated in the *N* × *N* matrix, *ppNEdgeTF*.

The last procedure in the algorithm is to integrate all the prior knowledge and compute the final post-inference network. The final post-inference network from the algorithm is the *N* ×*N* matrix *API* (acronym for Advanced post-inference), which contains the pairwise relation between all the genes considered. The *API* is obtained from the *infMat* by: a) removing the wrong interactions identified in the *ppNEdge*; b) adding the expected correct interactions in the *ppEdge*; c) adding the increased weights of inferred edges from transcription factors in the *ppEdgeTF*; d) removing the wrong edges from the non-transcription factors by scaling and subtracting the *ppNEdgeTG;* e) including the edge priors by scaling and adding the *edgePrior;* and f) removing the non-edge priors by scaling and subtracting the *nEdgePrior*. Scaling factor *w*, is chosen such that even if an edge or a non-edge is wrongly predicted in the post-inference procedures, they can be corrected by the prior knowledge. The pairwise interaction value for any two genes *i* and j, in the post-inference network matrix *API* is defined as,

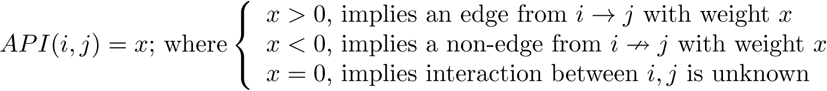

#### Theorem 1.

*The Advanced algorithm has a computational complexity of O(nN^2^* + N^3^).

*Proof*. The procedure to find the edge priors without data support, needs to check if *n* samples of the experimental data supports each of the edge priors (for *N* genes there can be a maximum of *N*^2^ edge priors). Thus, the complexity of this procedure is *O*(*nN*^2^).

The procedure to identify and remove the ancestor-FPs needs to consider each edge prior (maximum of *N*^2^) and the inferred parents for each of these edge priors (there can be a maximum of *N* parents). Thus, the complexity of this procedure is *O*(*N*^3^). Similarly, the complexity of the procedure to identify and remove the descendant-FPs is also *O*(*N^3^*).

The procedure to incorporate the knowledge of regulators consists of two steps. The first step of increasing the weight of inferred edges from transcription factors, identifies all the transcription factors (maximum of *N*) and the inferred target-genes for each of these transcription factors (maximum of *N*). Thus, the complexity is *O*(*N*^2^). The second step of removing non-edges from non-transcription factors identifies all the non-transcription factors (maximum of *N*) and then all its non-edges (maximum of *N*). Thus, the complexity is again *O*(*N*^2^). Therefore, the overall complexity of the procedure is *O*(*N*^2^).

The procedure to compute the final post-inference network needs to find the maximum value of the pairwise matrices as the scaling factor *w* (*O*(*N*^2^) complexity) and then compute the pairwise matrix *API* (*O*(*N*^2^) complexity). Thus, the complexity of this procedure is *O*(*N*^2^).

Therefore the overall complexity of the algorithm is

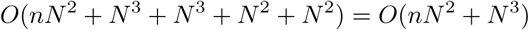

The complexity of the algorithm will be *O*(*nN*^2^+*N*^3^) for a fully connected network. For a typical GRN, the complexity will be much lower because the network will be sparse and the total number of edge priors will never be *N*^2^. The sparseness property can be taken advantage of by storing the edge priors in memory and not checking all the possible edge priors each time. In the algorithm implementation, the edge priors are stored in a list and read when required. Thus the complexity of considering all the edge priors is *O*(*N*) (assuming that the number of edge priors is proportional to the number of genes in the network). Further, due to the scale-free property of the GRNs (Albert, 2005; Barabasi and Oltvai, 2004), the number of parents for a gene will be much smaller than *N*. Due to these factors, the real complexity of the algorithm will be around *O*(*nN* + *N*^2^).

### 2.8 Metric to check data support for edge priors

The different reasons that can cause an edge prior interaction to lack support from the gene expression data are a) the interaction does not occur in the experimental conditions considered; b) the regulator and its target-gene mRNA transcripts are not correlated; c) the interaction is missed due to error or noise in the experimental data; or d) the prior knowledge is wrong. Thus, if an edge prior is not supported, it is an interesting candidate for further study.

There are many metrics to check if a prior edge is supported by the data such as: a) correlation between the genes, b) mutual information (MI) between the genes, or c) some other pairwise score or distance measure. Of all these, the MI is considered to be among the best because it can detect linear and non-linear dependencies, is well studied, and is reported to be better than other methods (Margolin et al., 2006). The data support with MI can be checked against a threshold (Margolin et al., 2006; Butte and Kohane, 2000) or with a statistical significance test (de Campos, 2006). The threshold value depends on the characteristics of a particular experimental data and needs to be recalibrated each time for new data. The statistical significance test does not have such drawbacks and is thus more intuitive. Pairwise MI with statistical significance test is used here because edge priors are direct pairwise relations. Though MI based tests have been reported before (de Campos, 2006; Wilczyński and Dojer, 2009; Vinh et al., 2011; Morshed et al., 2012), the mathematical derivation was not given and was cited to be in Kullback (1968). Kullback (1968) does not define the MI but uses relative entropy for all the results. Cover and Thomas (2006) (pp.19), presents a good definition of the MI and the relation between MI and relative entropy. Without the mathematical derivation, the details of computation such as the base of logarithm to be used and the assumptions involved are unclear. Thus, the MI statistical test derivation is provided here from the first principles.

#### Lemma 1.

*For any two variables* 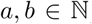, *the following approximation log_e_*, 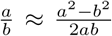*holds for* 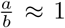 (Kullback (1968), pp-114).

#### Theorem 2.

*In pairwise statistical test of significance using MI, two genes x and y are said to be independent if* 2*NI*(*x*,*y*) *≤* χ^2^(α, *df*).

*Proof*. 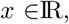 be the hypothesis that *x* and *y* are dependent with probability density *P*1 = *p*(*x*,*y*); and *H2* the null hypothesis that *x* and *y* are independent with probability density *P*2 = *p*(*x*)*p*(*y*). Then the mean information for discrimination in favour of H1 against H2 is given by (Kullback (1968) pp-8),

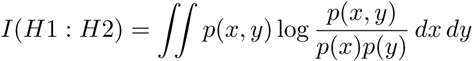

If *d_x_* and *d_y_* are the number of categories or classes in the population of *x* and *y* respectively, then

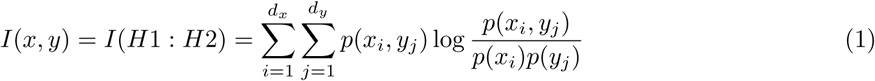
where, *I*(*x*,*y*) is known as the mutual information (Cover and Thomas (2006) pp-20) between the genes *x* and y; *p*(*x_i_*) and *p*(*y_j_*) are their individual probability distributions; and *p*(*x_i_, y*_j_) is their joint probability distribution. Thus, if *I*(*x, y*) = 0, then *x* and *y* are independent and larger values of mutual information implies higher dependence between the variables. However, even when the genes are independent, *I*(*x, y*) ≠ 0 can occur by chance. Thus, a statistical test of independence needs to be performed to identify the dependency between the variables.

The equation 1 in terms of H1 and H2 can be written as

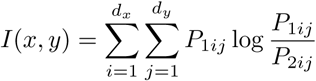

Applying the result of lemma 1, to the previous equation and deriving

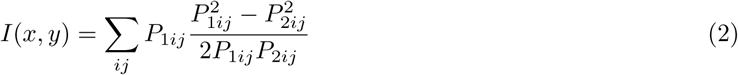

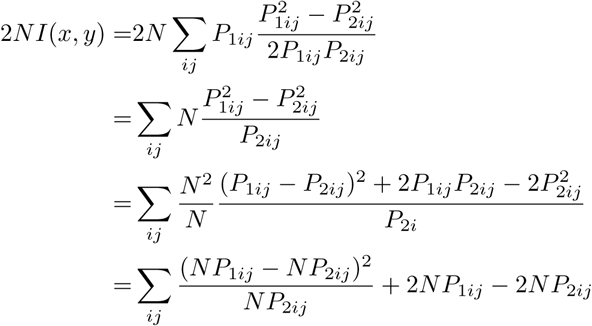

For the independence test, assume that the null hypothesis H2 (the variables are independent) is true. So the expected observation is *E_ij_* = *NP_2ij_* and the actual observation is *O_ij_* = *NP*_1*ij*_. Thus, the previous equation can be written as,

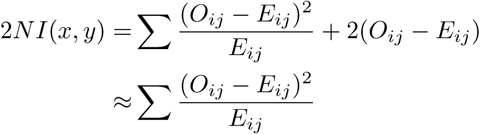

The right side of the equation is the form of the chi-squared test of independence with degrees of freedom *df* = (*d_x_* − 1)(d_y_ − 1), at significance level α. Thus,

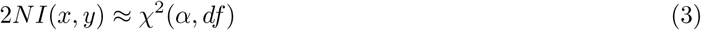

It is important to note that the above relation holds only when the MI is computed using *log_e_* because the equation 2 is valid only for logarithm with base *e*. Further, the equation 3 is valid when the null hypothesis is true (a similar result was stated for a two-way table with a slightly complicated derivation in Kullback (1968), pp-158). So, if *2NI*(*x*,*y*) > χ^2^(α, *df*), the null hypothesis is rejected at significance level α and the genes are considered dependent; else, if *2NI*(*x, y*) ≤ χ^2^(α, *df*), we fail to reject the null hypothesis at α and the genes are considered independent.

The χ^2^(α, *df*) is not dependent on the genes and their relation but represents the statistical chance of a particular value to appear for a system with *df* degrees of freedom at α significance. The 2*NI*(*x, y*) measures the strength of the dependence between genes *x* and *y* from the given experimental data. A higher dependence will cause a larger *2NI*(*x, y*). Thus, the above metric is also intuitive. During the significance test, *2NI*(*x, y*) is computed using the equation 1 from the experimental data and *χ^2^*(α, *df*) is obtained from the *χ^2^* distribution.

## 3 Experiment

This section presents the details of the networks and the data used for the study; the GRN inference method selected for the inference; the procedure followed to quantify the wrong edges predicted by the inference method; the performance metrics used to measure the improvement in GRN inference after the priors were incorporated; the prior knowledge used; and the implementation of the two novel post-inference prior algorithms — Basic and Advanced algorithms.

### 3.1 Networks and data

The DREAM5 transcription network inference challenge (Marbach et al., 2012) is the latest community-wide benchmarking study that compared the GRN inference performance of 35 inference methods. These 35 methods belonged to all the major categories of the GRN inference methods. The benchmarking study was performed on the gene expression data from the eukaryotic model organism — *S. cerevisiae*, the prokaryotic model organism — *E. coli*, the human pathogen — *Staphylococcus aureus*, and a realistic in silico network. These networks were named Network 4, 3, 2, and 1, respectively in the study. We selected the networks from this benchmark study because all the major GRN inference methods were compared; both biological and in silico networks were considered; the results and the original data are in public domain; and later studies have also used their networks and data. Further, because the DREAM5 results do not report *S. aureus* (named Network 2), this network is not considered here. The other three networks, Network 1, 3, and 4 are used here with the names Net1, Net3, and Net4, respectively. The details of these networks are as follows.

1. Realistic network (Net1): the DREAM5 in silico network. This network uses the known *E. coli* regulatory network with the gene expression data generated in silico but based on biological principles, using GeneNetWeaver (Schaffter et al., 2011) (version 4.0). There are 1643 genes and 195 transcription factors. The gold standard has 278,392 interactions, of which 4012 are edge priors and the rest 274,380 are non-edge priors.
2. *E. coli* network (Net3): the DREAM5 biological *E. coli* network. This network uses the currently known *E. coli* regulatory network with experimental gene expression data. There are 4511 genes and 334 known transcription factors. The gold standard has 152,280 interactions of which 2066 are edge priors and the rest 150,214 are non-edge priors.
3. Yeast network (Net4): the DREAM5 biological *S. cerevisiae* network. This network uses the currently known *S. cerevisiae* regulatory network with experimental gene expression data. There are 5950 genes and 333 known transcription factors. The gold standard has 227,202 interactions of which 3940 are edge priors and the rest 223,262 are non-edge priors.

For all the three networks, there are 805 lines of experimental data from 107 experiments which included 73 static and 34 time-series experiments. Thus, the gene expression data, prior knowledge (edge priors, non-edge priors, known transcription factors), and actual networks required for the study are obtained.

### 3.2 Selection of a GRN inference method

In the prior after inference methods proposed here, first an inferred network is obtained with a GRN inference method of user’s choice and then the prior knowledge is incorporated into this inferred network, in a post-inference step. Even though any GRN inference method can be used, we selected the GENIE3 (Huynh-Thu et al., 2010) method because it was a) the best-performer in DREAM5 Network inference challenge (Marbach et al., 2012); b) the best-performer in DREAM4 in silico network inference sub-challenge (Huynh-Thu et al., 2010); and c) based on an inference method that is not known to incorporate prior knowledge. Thus, GENIE3 being a high performing method which is not known to inherently incorporate the prior knowledge, is among the best candidates for a post-inference prior knowledge incorporation. GENIE3 will henceforth be referred to as Other1, which is its name in the DREAM5 results.

The inferred networks by the Other1 method for DREAM5 networks are also available in public domain. The inferred networks in DREAM5 have 100,000 predicted interactions as per the requirement of the DREAM5 competition. Each of the 100,000 predicted interaction has a confidence score associated with it, where a higher confidence score represents higher possibility of interaction. Thus, the results of the Other1 method on the DREAM5 networks are used as the inferred networks for this study.

### 3.3 Quantifying the wrong edge sources

It was shown in section 2.3 that the wrong edges in the inferred networks can be classified as originating from the strangers or one of the 13 relatives. This classification is applied to the inferred networks from the Other1 method to quantify the sources of wrong edges.

In the classification procedure, the known correct network and the inferred network for each of the three selected networks, Net1, 3, and 4, are obtained. Then, the quantification of the wrong edge sources in the inferred network can be performed by the following steps. For each gene in the correct network, all its parents, the 13 relatives, and the strangers are identified. Then, using the correct network, the wrong edges in the inferred network for each gene are identified. From these wrong edges, the wrong inferred parents for each gene are identified. Now, each of the wrong inferred parents is checked to determine whether it belongs to one of the 13 relatives or to the strangers that were identified earlier, for the gene. Thus, each wrong inferred parent can be classified. While checking if the wrong inferred parent is from any of the 13 relatives, the ancestors are checked first and then the descendants. Further, the different ancestor and descendant relatives are checked in the order given in Table 2.

This procedure to identify and classify the wrong edges is performed for all the genes in the inferred network. From this result, the total number of wrong edges, wrong edges due to strangers, and wrong edges from each of the 13 relatives can be quantified.

### 3.4 Performance metrics

The widely used performance metrics are applied to quantify the improvement given by the algorithms. These are: sensitivity or recall=TP/(TP+FN), is the fraction of correct edges detected; specificity=TN/(TN+FP), measures the fraction of correct non-edges detected; precision or positive predictive value=TP/(TP+FP), represents the fraction of correct edges in total predicted edges; and F1-score=2*precision*recall/(precision+recall), is the harmonic mean of precision and recall which is used to measure the performance in detecting true edges for skewed conditions. Here, TP is the true positives or the correct predicted edges; TN is the true negatives or the correct predicted non-edges; FP is the false positives (wrong edges), or the incorrect predicted edges; and FN is the false negatives or the incorrect predicted non-edges.

Further, standard metrics such as AUROC (area under ROC) and AUPR (area under precision-recall curve) are also computed using the script provided by the DREAM5. This will help to compare our results to the DREAM5 results. The AUROC measures the area under TPR (true positive rate) and FPR (false positive rate) curve where the TPR is the sensitivity and the FPR=FP/(FP+TN), is the fraction of non-edges wrongly classified as edges. For a random prediction, the value of the AUROC is 0.5 and in an ideal all correct prediction, it equals 1. The AUPR measures the area under precision-recall curve. For a random prediction, the AUPR is close to zero and for an ideal all correct prediction, it equals 1.

### 3.5 Basic post-inference algorithm

The inferred networks for the Net1, Net3, and Net4, obtained from the Other1 method are used to create the inferred network matrix, *infMat*. The prior knowledge matrices *edgePrior* and *nEdgePrior* are created from the ‘gold standards’ available for each network in the DREAM5 data. The prior knowledge is incorporated into the *infMat* according to the Algorithm 1. The post-inference matrix thus obtained is referred as the *BPI*. As discussed in section 2.6, the *BPI* is a weighted matrix with *BPI*(*i*,*j*) = *x*, where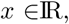, is the weight of the interaction from gene *i* to *j*. The weights are used to rank the edges while computing the AUROC and the AUPR. A threshold of *x* > 0 is applied to obtain a non-weighted matrix of edges to compute the other performance metrics such as precision, sensitivity, specificity, and F1-score.

The experiments are performed at different percentages of the total edge priors such as 0% (corresponding to the normal condition of having no priors), 10%, 25%, 50%, 75%, and 100% (all the priors used). For the 10%, 25%, 50%, and 75% experiments, the edge priors are chosen at random from all possible priors; and the experiments are repeated 10 times, to obtain the sample mean and standard deviation of the performance metrics. Further, separate experiments are conducted with all the non-edges along with the different percentage of edge priors to study the effect of the non-edge priors. Since only a small percentage of the non-edges are known for the networks studied here, different percentages of the non-edge priors are not considered.

### 3.6 Advanced post-inference algorithm

The *infMat, edgePrior*, and *nEdgePrior* variables are obtained using the same procedure described for the Basic algorithm, in section 3.5. The known transcription factors are available from the ‘input data’ of the DREAM5, from which the *tfPossible* is constructed. The gene expression data for each network is also available from the ‘input data’ in two files ‘expression_data’ and ‘chip_features’. These files are processed to separate the static and time-series gene expression data; and the data from different experiments. Once these inputs are obtained, the rest of the steps are given in Algorithm 2. Similar to the *BPI*, the *API* is also a weighted matrix with *API*(*i*,*j*) = *x*, where 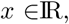 is the weight of the interaction from gene *i* to j. The AUROC and AUPR are computed by obtaining a ranked list of interactions based on this weight. A threshold of *x* > 0 is applied to obtain a non-weighted matrix of edges, to compute the other performance metrics.

Similar to the Basic algorithm, the experiments are performed for 0%, 10%, 25%, 50%, 75%, and 100% of the edge priors. Except for 0 and 100%, the priors are chosen at random from all possible priors and the experiments are repeated 10 times. Further, separate experiments are conducted with all the available non-edge priors along with the different percentage of edge priors.

The edge priors are checked for the data support with the MI statistical significance test. The edge priors are checked for support from all the available static experiment data samples, at α = 0.9. For the edge priors not supported in this condition, in the second stage, the support is checked with each static experiments individually. Since, individual experiments contain only lesser data samples, α = 0.999 is used. If any edge prior is still not supported, then in the third stage, the support from the individual time-series experimental data is checked for these edges, at α = 0.999. An edge that is not supported in all the three stages only will be reported as an edge prior without data support.

The following parameters are used to compute weights of edges and non-edges in the post-inference matrices *ppEdge, ppNEdge, ppEdgeTF*, and *ppNEdgeTF*: *weightP rior* = 2; *weightInfer* = 0.5; *weightDirect* = 0.5; *weightIndirect* = 0.25. The *weightP rior* is the weight of a known prior interaction such as an edge or a non-edge prior. It is given a weight of 2 because the maximum weight of an inferred interaction in the *infMat* is 1 or a -1. The prior interaction should have twice the weight of an inferred interaction so that even a wrong inferred interaction can be corrected. The *weightInfer* is the weight for an interaction, in the inferred network or supported by the data for a pairwise statistical significance test. Its value is lesser than the maximum weight of an edge in the *infMat* because the GRN inference methods generally use a more sophisticated technique than a simple pairwise test. The *weightDirect* is the weight given to a direct relation that is very likely to exist such as the inferred edges of a known transcription factor. The value of its weight is kept half of the maximum weight of an edge in the *infMat*. The *weightIndirect* is used when an indirect interaction is expected. For example if *TF*1 → *TF*2 → *TG* exists then it is likely that *TG* → *TF*1 does not exist. Therefore, this non-edge can be updated with *weightIndirect*. It needed a weight that is less than the *weightDirect* but greater than zero, so 0.25 is chosen.

## 4 Results and discussion

### 4.1 Quantifying the wrong edge sources

The wrong edges or the false positives in the inferred networks of the three selected networks, are shown in Table 3. The total false positives, the number of false positives from the strangers, relatives, and the 13 different relatives are given in the table. It can be seen from the table that all the networks predict a large number of total false positives. This is because the DREAM5 required each inferred network to have 100,000 interactions, which is much larger than the number of known correct interactions; resulting in a large number of false positives. Among the relatives, the non-descendant-FPs are much larger than the descendant-FPs. Further, more false positives originate from immediate ancestors than the ancestors farther away.

**Table 3:** Quantification of the total number of wrong edges and the different classes of wrong edges such as strangers and relatives, from the inferred networks of Net1, Net3, and Net4. The relatives are further classified into the 13 different relatives that are up to four-node distance. The different relatives are listed in the descending order of their frequency of occurrence in the inferred network.

| Class | Net1 | Net3 | Net4 | Distance |
| --- | --- | --- | --- | --- |
| Total | 96848 | 99386 | 99668 | - |
| Strangers | 56678 | 96536 | 93468 | - |
| Relatives | 40170 | 2850 | 6200 | - |
| Sib | 16545 | 1429 | 1985 | 2 |
| Aun | 9759 | 530 | 1195 | 3 |
| Nie | 3883 | 280 | 873 | 3 |
| Cus | 2768 | 194 | 651 | 4 |
| NCd | 2238 | 166 | 652 | 4 |
| GpS | 2095 | 89 | 325 | 4 |
| Gp | 1425 | 91 | 174 | 2 |
| GGp | 882 | 36 | 145 | 3 |
| GGGp | 376 | 8 | 116 | 4 |
| Chd | 150 | 25 | 39 | 1 |
| GGGc | 17 | 0 | 13 | 4 |
| Gc | 16 | 2 | 17 | 2 |
| GGc | 16 | 0 | 15 | 3 |

It was shown in section 2.3.1 that the false positives occur due to three reasons — indirect interactions, spurious interactions, and influence interactions. In Table 3, for the realistic network (Net1), 40% of the false positives are from the relatives, which are due to the indirect interactions. The rest 60% are from strangers which could be due to the spurious interactions caused by the noise and errors added in the data (Marbach et al., 2012). Further, a few stranger-FPs could occur due to the unclassified relatives at five-node distance and higher. The influence interactions due to protein-protein interactions and metabolites are not expected due to the in silico data generation method used (Schaffter et al., 2011).

In Table 3, for the *E. coli* (Net3) and yeast (Net4) real networks, the number of the relative-FPs are much lower compared to the Net1. There could be more than one reason for this occurrence. In real networks, the influence edges due to protein-protein interactions and metabolites could be present. The noise in the data and other experimental errors could also be higher because real experimental data is used. Further, these networks are not gold standard (Marbach et al., 2012) which means that our knowledge of these networks is not complete and more undiscovered interactions could be present. If more interactions are indeed present, then, many current stranger-FPs could actually be correct edges or relative-FPs.

The quantity of each relative-FP as a fraction of the total relative-FPs is shown in figure 3. The sibling-FP, at 40%, is the largest individual fraction of the total relative-FPs, for the Net1. In all the three networks, the sibling-FP is the largest source of wrong edges, followed by Aunt-FP, Niece-FP and so on. Thus, it can be seen that all the three networks have a similar distribution of the different relative-FPs with respect to the total relative-FP.

**Figure 3:**
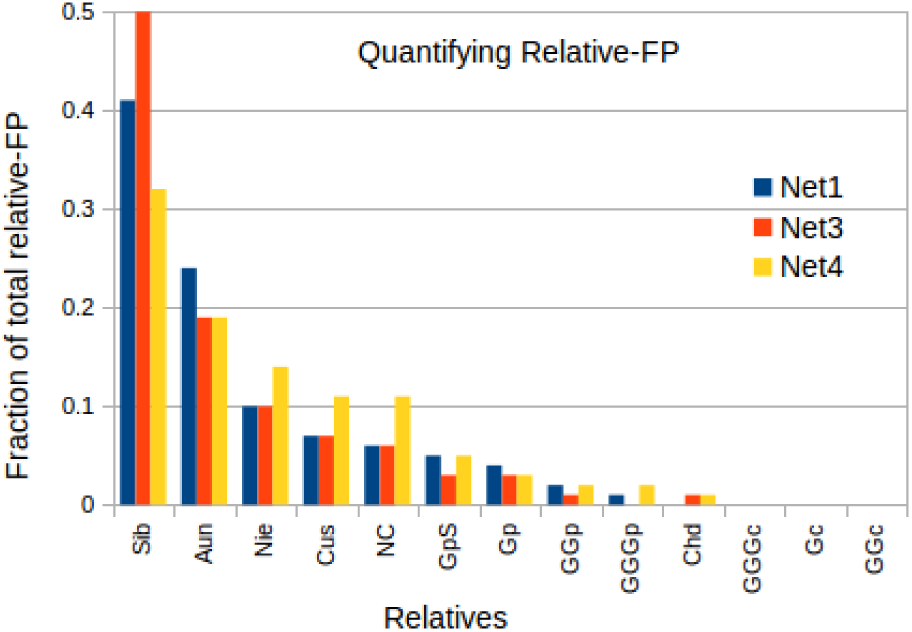
Fraction of wrong edges from each of the 13 relatives to the total relative-FP (false positives from all the relatives) for the three networks.

Thus, it was seen that all the inferred networks have a large number of relative-FPs caused due to the indirect interactions. The relative-FPs predominantly originate from the ancestor nodes than the descendants. Moreover, the nodes that are closer in node distance form a higher fraction of the relative-FPs. Further, all the three inferred networks show a similar distribution of the different relative-FPs. These observations validate the design and motivate the use of the Advanced post-inference algorithm.

### 4.2 Post-inference incorporation of priors

The results of the post-inference inclusion of the prior knowledge with the Basic and Advanced algorithms are presented for the three networks — Net1, Net3, and Net4. These results are compared to the results of a normal GRN inference without incorporating the prior knowledge (termed here as the Normal method), to identify the improvement. The study is performed only with the edge priors because a) the edge priors are the most widely used priors; b) it is difficult to obtain the non-edge priors; and c) only a small percentage of the non-edge priors are available for the networks considered here. However, because these algorithms can accept both edge and non-edge priors, the results with both priors are also presented.

#### 4.2.1 Net1: the realistic network

The performance of the Normal method, the Basic algorithm, and the Advanced algorithm for Net1 is given in Table 4. Only the edge priors are considered here. The averaged values of the different performance metrics and their sample standard deviations, at different percentages of the edge priors, are listed. It can be seen from the table that both the Basic and Advanced algorithms show equal or generally, much better performance than the Normal method, in all the performance metrics.

**Table 4:** Performance metrics for Normal method, Basic post-inference (BPI), and Advanced post-inference (API) algorithms on Net1 for the different percentage of edge priors (Edge%). In the performance metrics, TP is true positives and FP is false positives. A standard deviation equals to zero implies that its value is less than 0.001.

| Edge% | Method | TP | FP | Precision | Recall | F1-score | Specificity | AUROC | AUPR |
| --- | --- | --- | --- | --- | --- | --- | --- | --- | --- |
| - | Normal | 3152.0 | 96848.0 | 0.032 | 0.786 | 0.061 | 0.964 | 0.815 | 0.291 |
| 10 | BPI | 3237.4 | 96848.0 | 0.032 | 0.807 | 0.062 | 0.964 | 0.833 | 0.388 |
|  | API | 3219.6 | 96149.1 | 0.032 | 0.802 | 0.062 | 0.964 | 0.841 | 0.417 |
| 25 | BPI | 3371.1 | 96848.0 | 0.034 | 0.840 | 0.065 | 0.964 | 0.862 | 0.510 |
|  | API | 3332.5 | 95377.5 | 0.034 | 0.831 | 0.065 | 0.965 | 0.861 | 0.518 |
| 50 | BPI | 3585.9 | 96848.0 | 0.036 | 0.894 | 0.069 | 0.964 | 0.908 | 0.686 |
|  | API | 3545.8 | 94374.6 | 0.036 | 0.884 | 0.070 | 0.965 | 0.904 | 0.688 |
| 75 | BPI | 3799.4 | 96848.0 | 0.038 | 0.947 | 0.073 | 0.964 | 0.954 | 0.849 |
|  | API | 3776.6 | 93637.2 | 0.039 | 0.941 | 0.074 | 0.965 | 0.951 | 0.848 |
| 100 | BPI | 4012.0 | 96848.0 | 0.040 | 1.000 | 0.077 | 0.964 | 1.000 | 1.000 |
|  | API | 4012.0 | 93235.0 | 0.041 | 1.000 | 0.079 | 0.965 | 1.000 | 1.000 |
| Sample standard deviation |  |  |  |  |  |  |  |  |  |
| 10 | BPI | 4.115 | 0.000 | 0 | 0.001 | 0 | 0 | 0.001 | 0.002 |
|  | API | 3.921 | 177.363 | 0 | 0.001 | 0 | 0 | 0.002 | 0.007 |
| 25 | BPI | 10.354 | 0.000 | 0 | 0.003 | 0 | 0 | 0.001 | 0.002 |
|  | API | 9.312 | 153.893 | 0 | 0.002 | 0 | 0 | 0.001 | 0.004 |
| 50 | BPI | 15.271 | 0.000 | 0 | 0.004 | 0 | 0 | 0.002 | 0.003 |
|  | API | 15.915 | 196.964 | 0 | 0.004 | 0 | 0 | 0.002 | 0.004 |
| 75 | BPI | 13.108 | 0.000 | 0 | 0.003 | 0 | 0 | 0.002 | 0.003 |
|  | API | 14.577 | 101.427 | 0 | 0.004 | 0 | 0 | 0.002 | 0.002 |
| 100 | BPI | 0.000 | 0.000 | 0 | 0.000 | 0 | 0 | 0.000 | 0.000 |
|  | API | 0.000 | 0.000 | 0 | 0.000 | 0 | 0 | 0.000 | 0.000 |

The Basic algorithm predicts higher true positives than the Normal method which helps to improve all the performance metrics, except the false positives metric. This algorithm considers only the incorporation of the prior knowledge of interactions and does not remove the wrong edges. Since only the edge priors are available here, the algorithm cannot reduce the false positives (wrong edges) when compared to the Normal method.

The Advanced algorithm is better than the Normal method in all the performance metrics, including the false positives metric. This algorithm incorporates the prior knowledge of interactions and also tries to remove the wrong edges due to the indirect interactions. Thus, only the Advanced algorithm shows a reduction in the false positives when compared to the Normal method or the Basic algorithm. However, the Advanced algorithm predicts a slightly lower number of true positives than the Basic algorithm. As discussed in section 2.7, the Advanced algorithm uses an approximate method to identify the wrong edges when the prior knowledge of interactions are insufficient. This can cause a few correct edges to be lost. For example, in Table 4, for 10% edge priors, the Advanced algorithm identifies and removes around 700 wrong interactions and looses less than 20 correct interactions, when compared to the Basic algorithm. Moreover, the Advanced algorithm has consistently the lowest false positives and the highest precision, F1-score, specificity, AUROC, and AUPR among the three methods.

The improvements in performance metrics (computed from Table 4) for the Basic and Advanced algorithms when compared to the Normal method, are given in Table 5. In the table, true positives and false positives show improvement in the absolute number of edges. All the other metrics show percentage improvement with respect to the Normal method. For the 10% edge priors, the Basic algorithm has the highest improvement for the true positives with 85.4 more correct edges on an average than the Normal method which results in 2.7% better recall. For the Advanced method, the corresponding values are slightly lower with 67.6 more true positives and 2.0% better recall. However, for all the other metrics the Advanced algorithm outperforms the Basic algorithm. For the 10% edge priors, the Advanced algorithm reduces the false positives by around 700 edges on an average, which means it identifies and removes around 700 indirect interactions. It also improves AUROC by 3.2% and AUPR by 43.3%. Thus, even though the improvement in true positives is slightly lower, the Advanced algorithm displays a better overall improvement compared to the Basic algorithm.

**Table 5:**
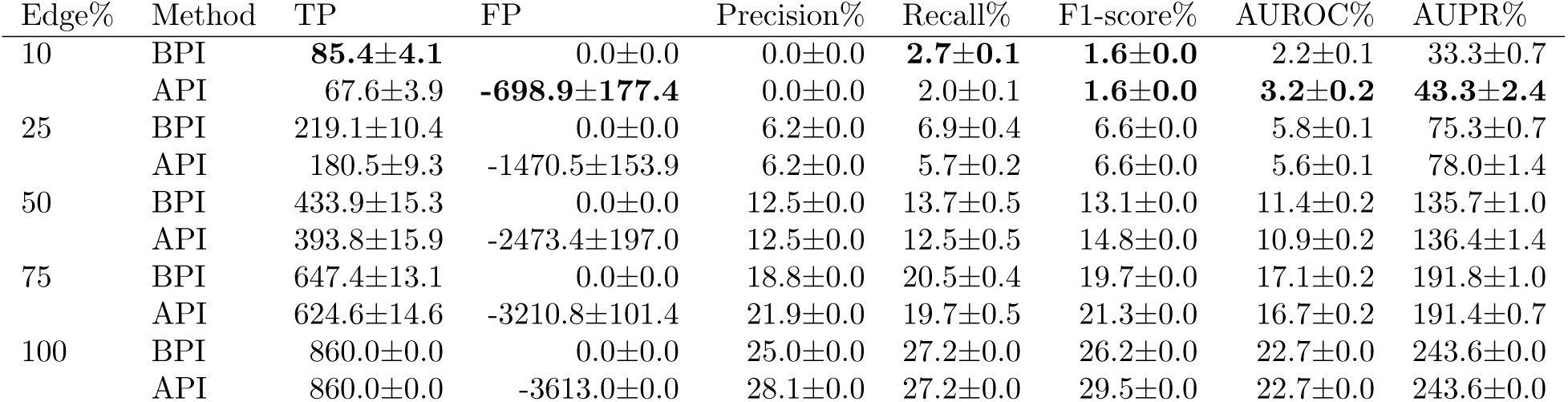
Improvement in performance metrics for Net1 with the Basic (BPI) and Advanced (API) algorithms when compared to the Normal method. The best-performance values for 10% edge priors are highlighted in bold typeface. In the performance metrics, TP is true positives and FP is false positives.

| Edge% | Method | TP | FP | Precision% | Recall% | F1-score% | AUROC% | AUPR% |
| --- | --- | --- | --- | --- | --- | --- | --- | --- |
| 10 | BPI | **85.4±4.1** | 0.0±0.0 | 0.0±0.0 | **2.7±0.1** | **1.6±0.0** | 2.2±0.1 | 33.3±0.7 |
|  | API | 67.6±3.9 | **-698.9±177.4** | 0.0±0.0 | 2.0±0.1 | **1.6±0.0** | **3.2±0.2** | **43.3±2.4** |
| 25 | BPI | 219.1±10.4 | 0.0±0.0 | 6.2±0.0 | 6.9±0.4 | 6.6±0.0 | 5.8±0.1 | 75.3±0.7 |
|  | API | 180.5±9.3 | -1470.5±153.9 | 6.2±0.0 | 5.7±0.2 | 6.6±0.0 | 5.6±0.1 | 78.0±1.4 |
| 50 | BPI | 433.9±15.3 | 0.0±0.0 | 12.5±0.0 | 13.7±0.5 | 13.1±0.0 | 11.4±0.2 | 135.7±1.0 |
|  | API | 393.8±15.9 | -2473.4±197.0 | 12.5±0.0 | 12.5±0.5 | 14.8±0.0 | 10.9±0.2 | 136.4±1.4 |
| 75 | BPI | 647.4±13.1 | 0.0±0.0 | 18.8±0.0 | 20.5±0.4 | 19.7±0.0 | 17.1±0.2 | 191.8±1.0 |
|  | API | 624.6±14.6 | -3210.8±101.4 | 21.9±0.0 | 19.7±0.5 | 21.3±0.0 | 16.7±0.2 | 191.4±0.7 |
| 100 | BPI | 860.0±0.0 | 0.0±0.0 | 25.0±0.0 | 27.2±0.0 | 26.2±0.0 | 22.7±0.0 | 243.6±0.0 |
|  | API | 860.0±0.0 | -3613.0±0.0 | 28.1±0.0 | 27.2±0.0 | 29.5±0.0 | 22.7±0.0 | 243.6±0.0 |

The performance of the Basic and the Advanced algorithms for the different performance metrics and edge prior percentages, can be compared using radar charts. In the radar charts here, the performance of the Normal method is assumed as the baseline of zero. Any increase of performance from the Normal method is scaled to [0,1] range, with 0 representing no improvement and 1 representing the highest improvement for that particular performance metric, among all the edge prior percentages in the Basic and the Advanced algorithms. Any decrease of performance is similarly scaled between [-1,0]. Thus, each performance metric is individually scaled between [-1,1], reflecting the increase or decrease in performance over the different edge prior percentages. Since the highest improvement selects the best-performance among the Basic and the Advanced algorithms, the radar chart can also compare the performance between the two algorithms.

The performance of the three algorithms (Normal, Basic, and Advanced) for the different performance metrics and edge prior percentages, can be seen in the radar charts in figure 4. Figure 4(a) shows the performance of the Normal and the Basic algorithms. The Basic algorithm shows higher performance in all the metrics, except false positives metric, when compared to the Normal method. Further, for the Basic algorithm, as the prior percentage increases, all the performance metrics (except false positive) also improve. Thus, 100% prior shows the best improvement, which is intuitive because all the correct edges are detected. However, precision and F1-score do not reach the maximum value even at 100% priors because the Advanced algorithm shows better improvement.

**Figure 4:**
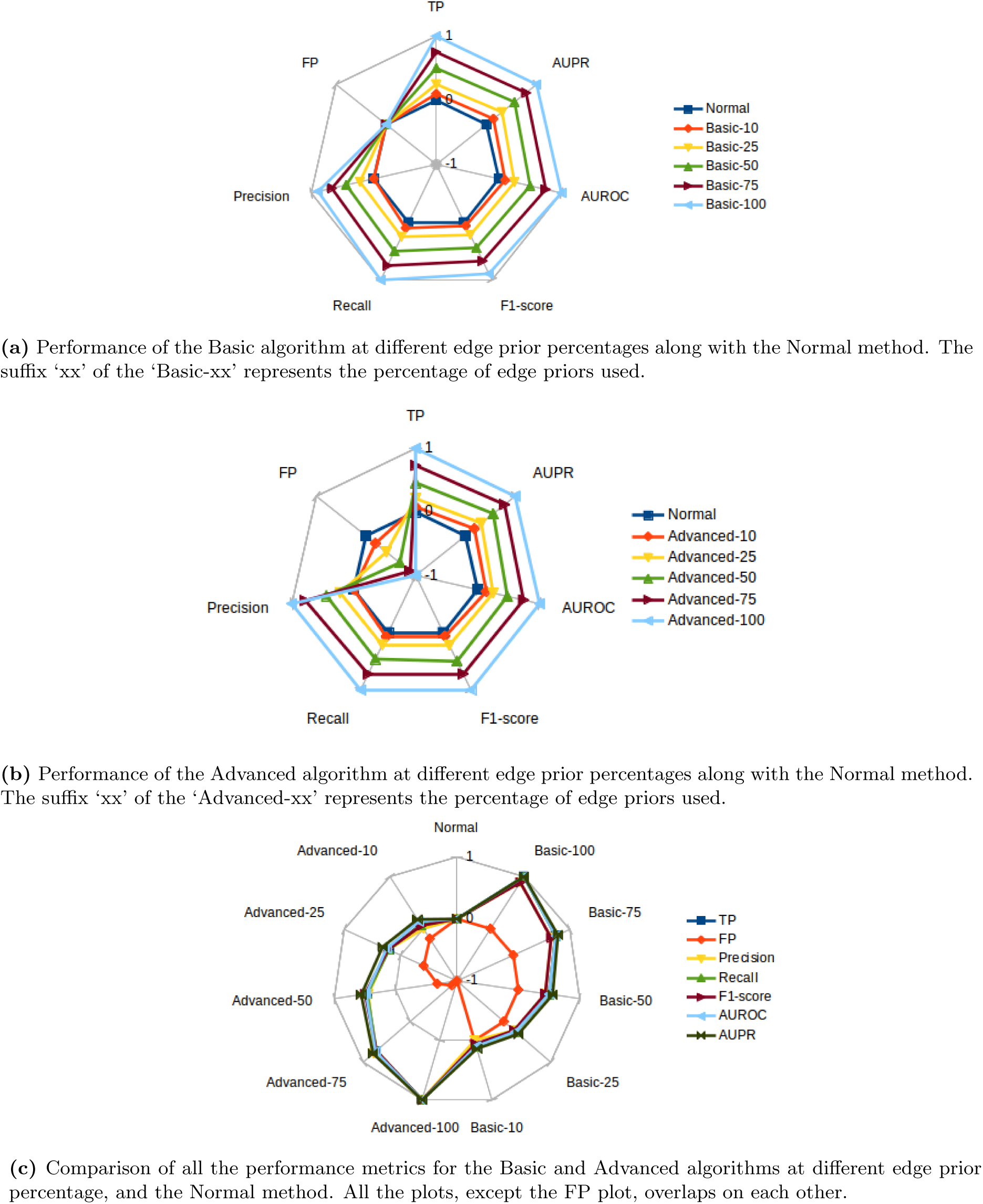
A Comparison of performances of Normal, Basic, and Advanced algorithms on Net1 for different performance metrics at 10%, 25%, 50%, 75%, and 100% edge priors. The performance of the Normal method is considered as the base performance at the value 0. A performance higher than Normal is in the range [0,1], and a performance lower than Normal is in the range [-1,0]. In the performance metrics, TP is true positives and FP is false positives.

Figure 4(b) shows the performance of the Normal and the Advanced algorithms for the different performance metrics. It can be seen that the Advanced algorithm shows improvement in all the performance metrics, including the false positives metric, when compared to the Normal method. As the prior percentage increases, all the performance metrics of the Advanced algorithm also show improvement.

Figure 4(c) shows the performance of the Normal, Basic, and Advanced algorithms in a single chart for the different performance metrics and prior percentages. This chart enables a comparison between the performance of the Basic and Advanced algorithms for the different metrics. From the symmetry of the plots in the chart, it can be seen that the Basic and Advanced algorithms have similar performances for all the performance metrics at the different edge prior percentages, except for the false positives metric. Further, among the Basic and the Advanced algorithms, only the Advanced algorithm shows a reduction in the false positives and thus, reduction in the wrong edges.

Thus, for the Net1, the Basic and Advanced algorithms show similar overall improvement when compared to the Normal method, for all the performance metrics except the wrong edges. Only the Advanced algorithm shows a reduction in the wrong edges and thus, shows an overall improvement across all the metrics.

#### 4.2.2 Net3: the E. coli network

The performance of the three algorithms for the Net3, are given in Table 6. Similar to the results of Net1, both the Basic and the Advanced algorithms show at least equal or generally, much better performance than the Normal method in all the performance metrics. The Basic algorithm predicts higher true positives than the Normal, which improves all the performance metrics except the false positives metric.

**Table 6:** Performance metrics of Normal method, Basic post-inference (BPI), and Advanced post-inference (API) algorithms on Net3, for different percentage of edge priors (Edge%). In the performance metrics, TP is true positives and FP is false positives. A standard deviation equals to zero implies that its value is less than 0.001.

| Edge% | Method | TP | FP | Precision | Recall | F1-score | Specificity | AUROC | AUPR |
| --- | --- | --- | --- | --- | --- | --- | --- | --- | --- |
| - | Normal | 614.0 | 99386.0 | 0.006 | 0.297 | 0.012 | 0.995 | 0.617 | 0.093 |
| 10 | BPI | 762.4 | 99386.0 | 0.008 | 0.369 | 0.015 | 0.995 | 0.656 | 0.211 |
|  | API | 759.4 | 99298.4 | 0.008 | 0.368 | 0.015 | 0.995 | 0.656 | 0.196 |
| 25 | BPI | 979.0 | 99386.0 | 0.010 | 0.474 | 0.019 | 0.995 | 0.713 | 0.358 |
|  | API | 971.4 | 99112.5 | 0.010 | 0.470 | 0.019 | 0.995 | 0.712 | 0.355 |
| 50 | BPI | 1340.7 | 99386.0 | 0.013 | 0.649 | 0.026 | 0.995 | 0.809 | 0.583 |
|  | API | 1333.1 | 98837.9 | 0.013 | 0.645 | 0.026 | 0.995 | 0.807 | 0.581 |
| 75 | BPI | 1702.2 | 99386.0 | 0.017 | 0.824 | 0.033 | 0.995 | 0.904 | 0.794 |
|  | API | 1697.6 | 98630.6 | 0.017 | 0.822 | 0.033 | 0.995 | 0.903 | 0.793 |
| 100 | BPI | 2066.0 | 99386.0 | 0.020 | 1.000 | 0.040 | 0.995 | 1.000 | 1.000 |
|  | API | 2066.0 | 98459.0 | 0.021 | 1.000 | 0.040 | 0.995 | 1.000 | 1.000 |
| Sample standard deviation |  |  |  |  |  |  |  |  |  |
| 10 | BPI | 7.792 | 0.000 | 0 | 0.004 | 0 | 0 | 0.002 | 0.002 |
|  | API | 8.656 | 36.713 | 0 | 0.004 | 0 | 0 | 0.002 | 0.005 |
| 25 | BPI | 10.066 | 0.000 | 0 | 0.005 | 0 | 0 | 0.003 | 0.003 |
|  | API | 9.732 | 34.375 | 0 | 0.005 | 0 | 0 | 0.002 | 0.003 |
| 50 | BPI | 6.945 | 0.000 | 0 | 0.003 | 0 | 0 | 0.002 | 0.002 |
|  | API | 7.824 | 57.398 | 0 | 0.004 | 0 | 0 | 0.002 | 0.002 |
| 75 | BPI | 13.164 | 0.000 | 0 | 0.006 | 0 | 0 | 0.003 | 0.003 |
|  | API | 12.963 | 44.970 | 0 | 0.006 | 0 | 0 | 0.003 | 0.003 |
| 100 | BPI | 0.000 | 0.000 | 0 | 0.000 | 0 | 0 | 0.000 | 0.000 |
|  | API | 0.000 | 0.000 | 0 | 0.000 | 0 | 0 | 0.000 | 0.000 |

The Advanced algorithm is better than the Normal method in all the performance metrics, including the false positives metric. Only the Advanced algorithm shows a reduction in the false positives when compared to the Normal method or the Basic algorithm. However, the Advanced algorithm predicts a slightly lower number of true positives than the Basic algorithm, similar to the Net1 results. In Table 6, for 10% edge priors, the Advanced algorithm identifies and removes around 90 wrong interactions while loosing only around 3 correct interactions when compared to the Basic algorithm. Thus, the Advanced algorithm consistently shows good overall performance among the three methods.

The improvements in the performance metrics (computed from Table 6) of the Basic and the Advanced algorithms when compared to the Normal method are given in Table 7. In the table, the true positives and the false positives show improvement in absolute number of edges while the other metrics show percentage improvement. For the 10% edge priors, the Basic algorithm has the highest improvement for the true positives with 148.4 more correct edges on an average, than the Normal method, which results in 24.2% better recall. In the Advanced method the corresponding values are slightly lower at 145.4 improved true positives and 23.9% improved recall. However, for most of the other metrics the Advanced algorithm has comparable or better performance than the Basic algorithm. For the 10% edge priors, the Advanced algorithm reduces the false positives by around 90 edges, on an average. This means that it identifies and removes around 90 indirect interactions. Thus, even though the improvement in true positives is slightly lower than the Basic algorithm, only the Advanced algorithm shows an overall improvement in all the performance metrics.

**Table 7:** Improvement in performance metrics for Net3 with the Basic (BPI) and Advanced (API) algorithms when compared to the Normal method. The best-performance values for 10% edge priors are highlighted in bold typeface. In the performance metrics, TP is true positives and FP is false positives.

| Edge% | Method | TP | FP | Precision% | Recall% | F1-score% | AUROC% | AUPR% |
| --- | --- | --- | --- | --- | --- | --- | --- | --- |
| 10 | BPI | **148.4±7.8** | 0.0±0.0 | **33.3±0.0** | **24.2±1.4** | **25.0±0.0** | **6.3±0.3** | **126.9±2.1** |
|  | API | 145.4±8.7 | **-87.6±36.7** | **33.3±0.0** | 23.9±1.4 | **25.0±0.0** | **6.3±0.3** | 110.8±5.4 |
| 25 | BPI | 365.0±10.1 | 0.0±0.0 | 66.7±0.0 | 59.6±1.7 | 58.3±0.0 | 15.6±0.5 | 284.9±3.2 |
|  | API | 357.4±9.7 | -273.5±34.4 | 66.7±0.0 | 58.2±1.7 | 58.3±0.0 | 15.4±0.3 | 281.7±3.2 |
| 50 | BPI | 726.7±6.9 | 0.0±0.0 | 116.7±0.0 | 118.5±1.0 | 116.7±0.0 | 31.1±0.3 | 526.9±2.1 |
|  | API | 719.1±7.8 | -548.1±57.4 | 116.7±0.0 | 117.2±1.4 | 116.7±0.0 | 30.8±0.3 | 524.7±2.1 |
| 75 | BPI | 1088.2±13.2 | 0.0±0.0 | 183.3±0.0 | 177.4±2.0 | 175.0±0.0 | 46.5±0.5 | 753.8±3.2 |
|  | API | 1083.6±13.0 | -755.4±45.0 | 183.3±0.0 | 176.8±2.0 | 175.0±0.0 | 46.4±0.5 | 752.7±3.2 |
| 100 | BPI | 1452.0±0.0 | 0.0±0.0 | 233.3±0.0 | 236.7±0.0 | 233.3±0.0 | 62.1±0.0 | 975.3±0.0 |
|  | API | 1452.0±0.0 | -927.0±0.0 | 250.0±0.0 | 236.7±0.0 | 233.3±0.0 | 62.1±0.0 | 975.3±0.0 |

Similar to the Net1 results, the performance comparison of the three algorithms for the different performance metrics and edge prior percentages, for the Net3, is shown using radar charts in figure 5. Figure 5(a) shows the performance comparison of the Normal and the Basic algorithms. The Basic algorithm shows higher performance in all metrics, except the false positives metric. Further, as the prior percentage increases for the Basic algorithm, all the performance metrics (except false positives) improve and the 100% prior shows the best improvement.

**Figure 5:**
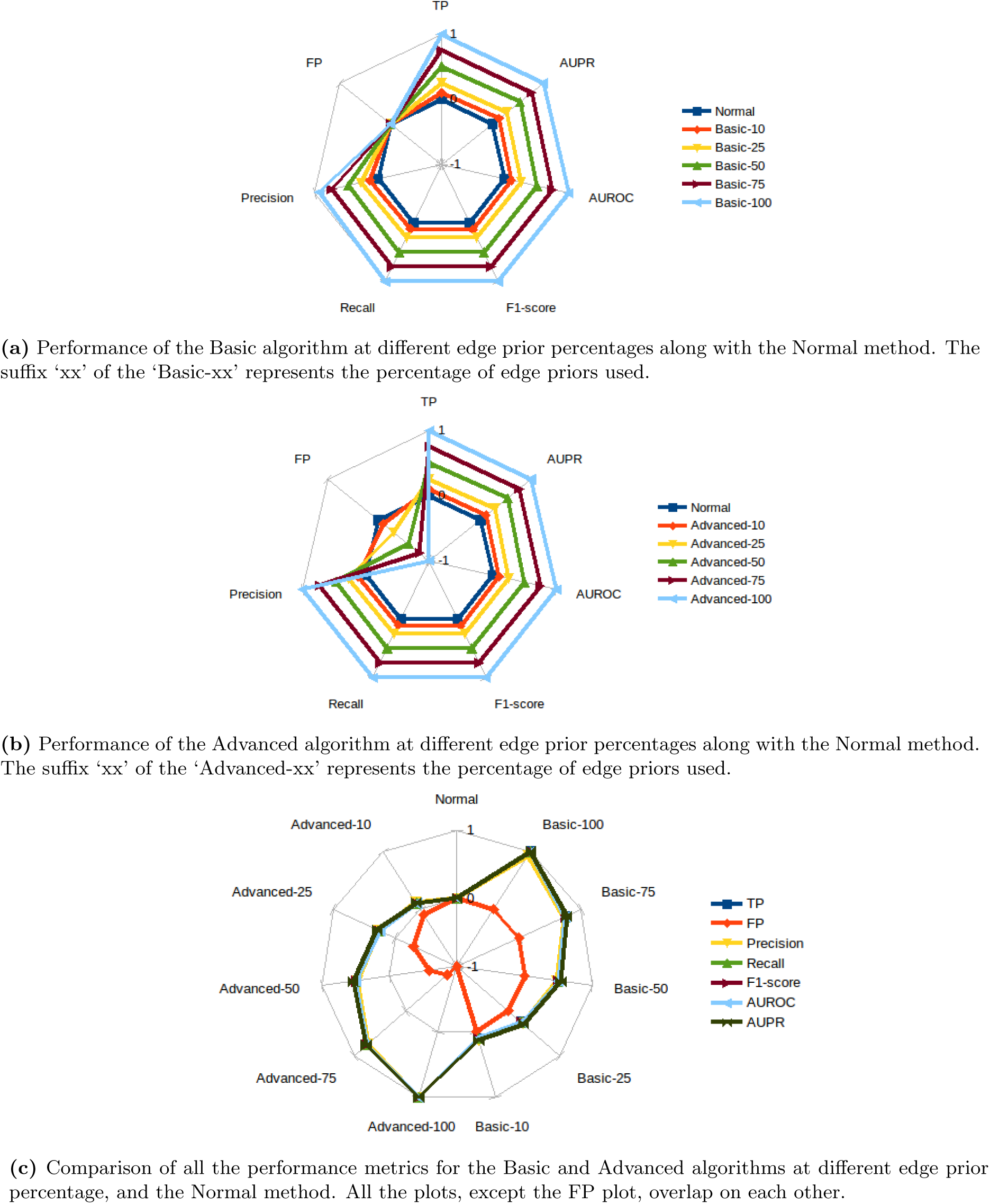
Comparative performance of Normal, Basic, and Advanced algorithms on Net3 for the different performance metrics at 0%, 25%, 50%, 75%, and 100% edge priors. The Normal method performance is considered as the base performance at the value 0. Performance higher than Normal is in the range [0,1], and performance lower than Normal is in the range [-1,0]. In the performance metrics, TP is true positives and FP is false positives.

Figure 5(b) shows the performance of the Normal and the Advanced algorithms. The Advanced algorithm shows improvement in all the performance metrics including the false positives, when compared to the Normal method. Figure 5(c) shows the performance of the Normal, the Basic, and the Advanced algorithms in a single chart. From the symmetry of the plots in the chart, it can be seen that the Basic and the Advanced algorithms show a similar performance for all the performance metrics across the different edge prior percentages, except for the false positives metric. Only the Advanced algorithm shows a reduction in the false positives and thus, a reduction in the wrong edges.

Thus, for the Net3 network, the Basic and Advanced algorithms show similar improvement when compared to the Normal method in all the performance metrics, except the wrong edges. Only the Advanced algorithm shows a reduction in the wrong edges and therefore, an overall improvement across all the performance metrics. These results are similar to the Net1 results.

#### 4.2.3 Net4: the yeast network

The performances of the three algorithms in Net4 are in Table 8. Similar to the results of Net1 and Net3, both the Basic and the Advanced algorithms have at least equal or generally, much better performance than the Normal method in all the performance metrics. The Basic algorithm is able to predict higher true positives than the Normal which improves all the other performance metrics, except the false positives.

**Table 8:** Performance metrics of Normal method, Basic post-inference (BPI), and Advanced post-inference (API) algorithms on Net4, for different percentage of edge priors (Edge%). In the performance metrics, TP is true positives and FP is false positives. A standard deviation equals to zero implies that its value is less than 0.001.

| Edge% | Method | TP | FP | Precision | Recall | F1-score | Specificity | AUROC | AUPR |
| --- | --- | --- | --- | --- | --- | --- | --- | --- | --- |
| - | Normal | 332.0 | 99668.0 | 0.003 | 0.084 | 0.006 | 0.997 | 0.518 | 0.021 |
| 10 | BPI | 693.0 | 99668.0 | 0.007 | 0.176 | 0.013 | 0.997 | 0.566 | 0.132 |
|  | API | 691.4 | 99499.5 | 0.007 | 0.175 | 0.013 | 0.997 | 0.566 | 0.136 |
| 25 | BPI | 1238.0 | 99668.0 | 0.012 | 0.314 | 0.024 | 0.997 | 0.639 | 0.287 |
|  | API | 1233.5 | 99246.3 | 0.012 | 0.313 | 0.024 | 0.997 | 0.639 | 0.290 |
| 50 | BPI | 2131.9 | 99668.0 | 0.021 | 0.541 | 0.040 | 0.997 | 0.758 | 0.532 |
|  | API | 2126.2 | 98858.0 | 0.021 | 0.540 | 0.041 | 0.997 | 0.758 | 0.533 |
| 75 | BPI | 3037.3 | 99668.0 | 0.030 | 0.771 | 0.057 | 0.997 | 0.879 | 0.770 |
|  | API | 3033.2 | 98623.0 | 0.030 | 0.770 | 0.057 | 0.997 | 0.879 | 0.770 |
| 100 | BPI | 3940.0 | 99668.0 | 0.038 | 1.000 | 0.073 | 0.997 | 1.000 | 1.000 |
|  | API | 3940.0 | 98415.0 | 0.038 | 1.000 | 0.074 | 0.997 | 1.000 | 1.000 |
| Sample standard deviation |  |  |  |  |  |  |  |  |  |
| 10 | BPI | 4.944 | 0.000 | 0 | 0.001 | 0 | 0 | 0.001 | 0.000 |
|  | API | 5.038 | 29.387 | 0 | 0.001 | 0 | 0 | 0.001 | 0.001 |
| 25 | BPI | 5.735 | 0.000 | 0 | 0.001 | 0 | 0 | 0.001 | 0.001 |
|  | API | 5.482 | 46.555 | 0 | 0.001 | 0 | 0 | 0.001 | 0.001 |
| 50 | BPI | 7.109 | 0.000 | 0 | 0.002 | 0 | 0 | 0.001 | 0.001 |
|  | API | 7.843 | 48.854 | 0 | 0.002 | 0 | 0 | 0.001 | 0.001 |
| 75 | BPI | 6.584 | 0.000 | 0 | 0.002 | 0 | 0 | 0.001 | 0.001 |
|  | API | 6.391 | 41.266 | 0 | 0.002 | 0 | 0 | 0.001 | 0.001 |
| 100 | BPI | 0 | 0 | 0 | 0 | 0 | 0 | 0 | 0 |
|  | API | 0 | 0 | 0 | 0 | 0 | 0 | 0 | 0 |

The Advanced algorithm is better than the Normal method in all the performance metrics, including the false positives. Thus, only the Advanced algorithm shows a reduction in the false positives when compared to the Normal method or the Basic algorithm. In Table 8, for the 10% edge priors, the Advanced algorithm identifies and removes around 170 wrong interactions while loosing only around 1 correct interaction when compared to the Basic algorithm. Thus, the Advanced method consistently shows the best overall performance among the three algorithms.

The improvements in the performance metrics (computed from Table 8) of the Basic and the Advanced algorithms when compared to the Normal method, are given in Table 9. For the 10% edge priors, the Basic algorithm has the highest improvement for the true positives with 361 more correct edges on an average, than the Normal method. This results in 109.5% better recall. For the Advanced method, the corresponding values are comparable at 359.4 more true positives and 108.3% improved recall. For other metrics also the Advanced algorithm has equal or better performance than the Basic algorithm. The Advanced algorithm reduces the false positives by around 170 edges which is the magnitude of the indirect interactions that are identified and removed. Thus, the Advanced algorithm shows the best overall improvement in all the performance metrics.

**Table 9:** Improvement in performance metrics for Net4 with the Basic (BPI) and Advanced (API) algorithms when compared to the Normal method. The best-performance values for 10% edge priors are highlighted in bold typeface. In the performance metrics, TP is true positives and FP is false positives.

| Edge% | Method | TP | FP | Precision% | Recall% | F1-score% | AUROC% | AUPR% |
| --- | --- | --- | --- | --- | --- | --- | --- | --- |
| 10 | BPI | **361.0±4.9** | 0.0±0.0 | **133.3±0.0** | **109.5±0.0** | **116.7±0.0** | **9.3±0.0** | 528.6±0.0 |
|  | API | 359.4±5.0 | **-168.5±29.4** | **133.3±0.0** | 108.3±0.0 | **116.7±0.0** | **9.3±0.0** | **547.6±0.0** |
| 25 | BPI | 906.0±5.7 | 0.0±0.0 | 300.0±0.0 | 273.8±0.0 | 300.0±0.0 | 23.4±0.0 | 1266.7±0.0 |
|  | API | 901.5±5.5 | -421.7±46.6 | 300.0±0.0 | 272.6±0.0 | 300.0±0.0 | 23.4±0.0 | 1281.0±0.0 |
| 50 | BPI | 1799.9±7.1 | 0.0±0.0 | 600.0±0.0 | 544.0±0.0 | 566.7±0.0 | 46.3±0.0 | 2433.3±0.0 |
|  | API | 1794.2±7.8 | -810.0±48.9 | 600.0±0.0 | 542.9±0.0 | 583.3±0.0 | 46.3±0.0 | 2438.1±0.0 |
| 75 | BPI | 2705.3±6.6 | 0.0±0.0 | 900.0±0.0 | 817.9±0.0 | 850.0±0.0 | 69.7±0.0 | 3566.7±0.0 |
|  | API | 2701.2±6.4 | -1045.0±41.3 | 900.0±0.0 | 816.7±0.0 | 850.0±0.0 | 69.7±0.0 | 3566.7±0.0 |
| 100 | BPI | 3608.0±0.0 | 0.0±0.0 | 1166.7±0.0 | 1090.5±0.0 | 1116.7±0.0 | 93.0±0.0 | 4661.9±0.0 |
|  | API | 3608.0±0.0 | -1253.0±0.0 | 1166.7±0.0 | 1090.5±0.0 | 1133.3±0.0 | 93.0±0.0 | 4661.9±0.0 |

Similar to the Net1 and Net3 results, the performances of the three algorithms for Net4 are compared using the radar charts in figure 6. Figure 6(a) compares the performance of the Normal and the Basic algorithms. The Basic algorithm shows better performance in all the metrics, except for the false positives. Further, as the prior percentage increases for the Basic algorithm, all the performance metrics (except the false positives) improve and the 100% prior shows the best improvement. Figure 6(b) compares the performance of the Normal and the Advanced algorithms. The Advanced algorithm shows improvement for all the performance metrics, including the false positives metric, when compared to the Normal method. Figure 6(c) compares the performance of the Normal, the Basic, and the Advanced algorithms in a single chart. The symmetry in the plots and overlapping plots show that the Basic and the Advanced algorithms have similar performances in all the performance metrics at the same edge prior percentages, except for the false positives metric. Only the Advanced algorithm shows a reduction in the false positives and thus, achieves reduction in the wrong edges.

Thus, for Net4, the Basic and the Advanced algorithms show similar improvements when compared to the Normal method for all the performance metrics except the wrong edges. Only the Advanced algorithm shows a reduction in wrong edges and therefore, an overall improvement across all the performance metrics. These results are similar to the results from Net1 and Net3.

**Figure 6:**
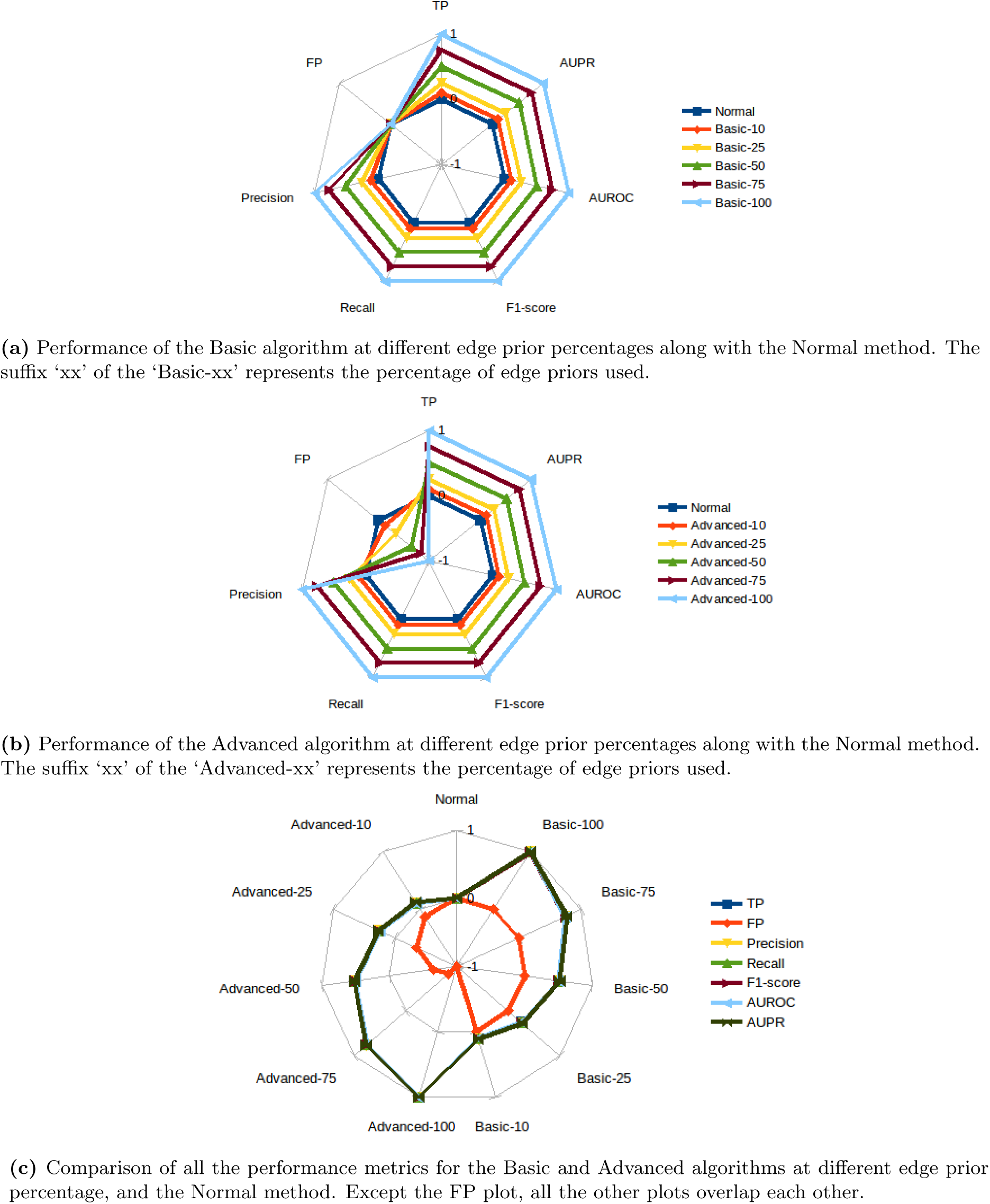
Comparative performance of Normal, Basic, and Advanced algorithms on Net4 for the different performance metrics at 10%, 25%, 50%, 75%, and 100% edge priors. The Normal method performance is considered as the base performance at the value 0. Performance higher than Normal is in the range [0,1], and performance lower than Normal is in the range [-1,0]. In the performance metrics, TP is true positives and FP is false positives.

#### 4.2.4 Comparison of Basic and Advanced algorithm

Figure 7 shows the improvements in correct edges (true positives) and reduction in wrong edges (false positives) given by the Basic and Advanced algorithms when compared to the Normal method in the three networks, Net1, Net3, and Net4. The improvements are shown for the 10%, 25%, 50%, 75%, and 100% of edge priors. For example, figure 7(a) shows that in Net1, for 10% edge priors, the Basic algorithm predicts around 100 correct edges more than the Normal method but figure 7(b) shows that the Basic algorithm exhibits no comparative reduction in the wrong edges. The Advanced algorithm, for the same conditions, predicts around 100 correct edges higher (figure 7(a)) and around 700 wrong edges lower (figure 7(b)) than the Normal method. The Advanced algorithm predicts slightly less number of correct edges than the Basic algorithm but only the Advanced algorithm shows reduction in wrong edges. This pattern can be seen for all the edge prior quantities and for all the three networks in figure 7. It can be seen from the figure that for the three networks at half the quantity (50%) of edge priors, on an average, the Basic algorithm predicts around 980 correct edges more than the Normal method while giving no reduction in the number of wrong edges. In contrast, the Advanced algorithm for the same conditions predicts around 970 correct edges more and removes around 1300 wrong edges when compared to the Normal method.

**Figure 7:**
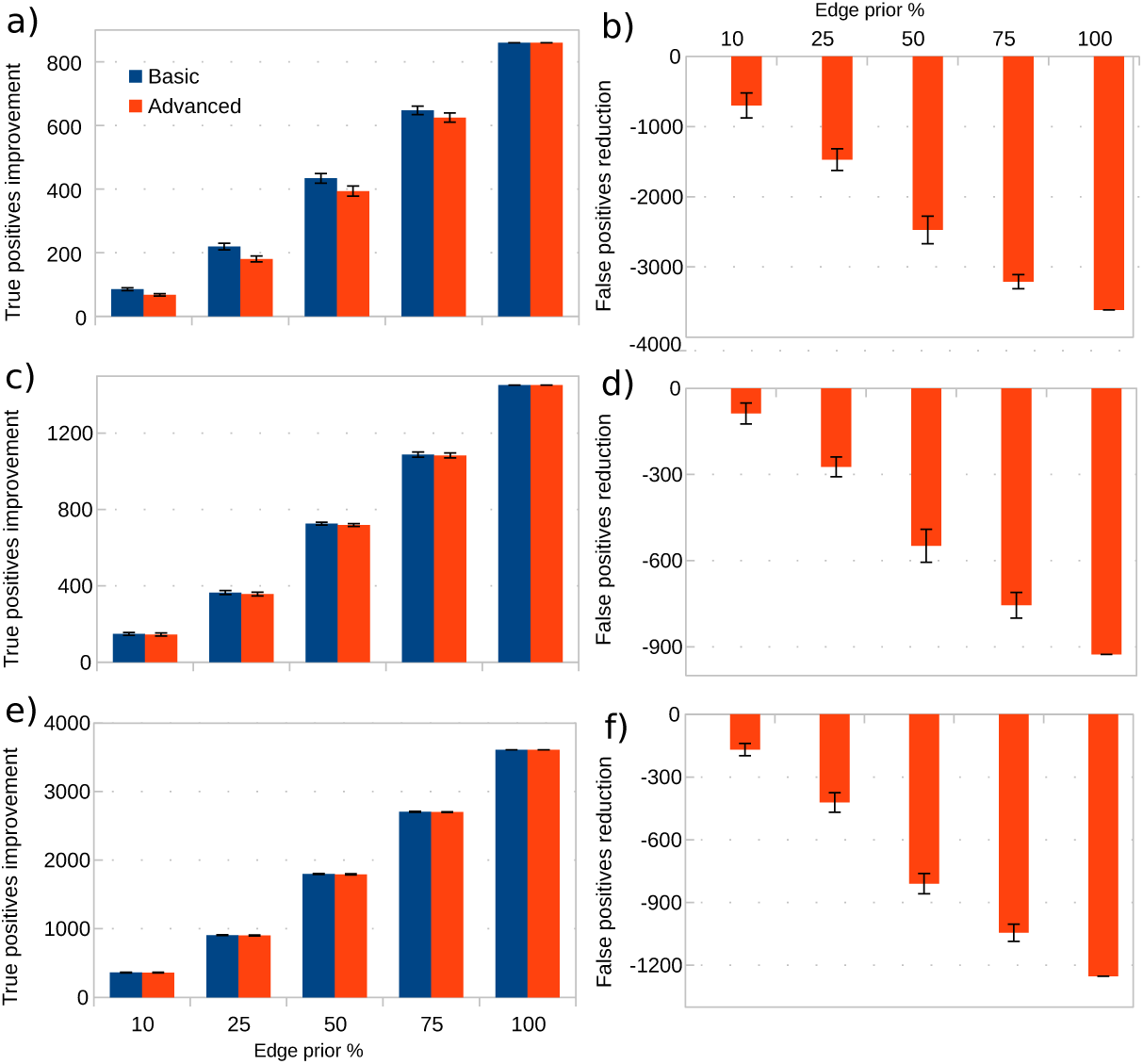
Improvement in true positives and reduction in false positives for the three networks with the Basic and Advanced algorithms when compared to the Normal method, at different edge prior percentages. a), c), and e) show the improvement in true positives for the Net1, Net3, and Net4, respectively. b), d), and f) show the reduction in false positives for the Net1, Net3, and Net4, respectively. Only the Advanced algorithm shows reduction in false positives and the Basic algorithm shows no reduction in false positives compared to the Normal method.

#### 4.2.5 Incorporation of edge and non-edge priors

The results presented previously on the Net1, Net3, and Net4 incorporated only the edge priors. However, the post-inference prior algorithms developed here can incorporate both the edge and the non-edge priors. Net1 is selected for this study because it is the only gold standard network in which all the edge priors are known. For the *E. coli* (Net3) and the yeast (Net4) networks all the edge priors may not be known (Marbach et al., 2012) and consequently, the classification of the wrong edges could be incorrect. Thus, the wrong-edge-removal procedure may not be quantified correctly.

As discussed in section 3.1, the Net1 has 1643 genes which can have 2,699,449 interactions (1643×1643). Since there are 4012 edge priors, the total possible non-edges are 2,699,449-4012=2,695,437. Out of the more than two-million non-edges, only around one-tenth (274,380) are available as priors in the DREAM5 data. Due to the low quantity of the non-edge priors known, all the available non-edge priors are always incorporated in the Basic and the Advanced algorithms; and only the edge prior quantity is varied between 0–100%. The results of this study are in Table 10.

**Table 10:** Performance metrics of Normal method, Basic post-inference (BPI), and Advanced post-inference (API) algorithms on Net1 with edge and non-edge priors. All the available non-edge priors are incorporated in the Basic and Advanced algorithms and only the quantity of the edge priors (Edge%) are varied. In the performance metrics, TP is true positives and FP is false positives. A standard deviation equals to zero implies that its value is less than 0.001.

| Edge% | Method | TP | FP | Precision | Recall | F1-score | Specificity | AUROC | AUPR |
| --- | --- | --- | --- | --- | --- | --- | --- | --- | --- |
| - | Normal | 3152.0 | 96848.0 | 0.032 | 0.786 | 0.061 | 0.964 | 0.815 | 0.291 |
| 10 | BPI | 3236.3 | 12083.0 | 0.211 | 0.807 | 0.335 | 0.996 | 0.903 | 0.817 |
|  | API | 3194.1 | 12017.1 | 0.210 | 0.796 | 0.332 | 0.996 | 0.898 | 0.807 |
| 25 | BPI | 3370.2 | 12083.0 | 0.218 | 0.840 | 0.346 | 0.996 | 0.920 | 0.849 |
|  | API | 3318.3 | 11908.3 | 0.218 | 0.827 | 0.345 | 0.996 | 0.914 | 0.837 |
| 50 | BPI | 3579.5 | 12083.0 | 0.229 | 0.892 | 0.364 | 0.996 | 0.946 | 0.898 |
|  | API | 3534.2 | 11784.6 | 0.231 | 0.881 | 0.366 | 0.996 | 0.940 | 0.888 |
| 75 | BPI | 3802.4 | 12083.0 | 0.239 | 0.948 | 0.382 | 0.996 | 0.974 | 0.951 |
|  | API | 3772.3 | 11699.2 | 0.244 | 0.940 | 0.387 | 0.996 | 0.970 | 0.944 |
| 100 | BPI | 4012.0 | 12083.0 | 0.249 | 1.000 | 0.399 | 0.996 | 1.000 | 1.000 |
|  | API | 4012.0 | 11628.0 | 0.257 | 1.000 | 0.408 | 0.996 | 1.000 | 1.000 |
| Sample standard deviation |  |  |  |  |  |  |  |  |  |
| 10 | BPI | 7.072 | 0.000 | 0.000 | 0.002 | 0.001 | 0 | 0.001 | 0.002 |
|  | API | 9.291 | 12.775 | 0.001 | 0.002 | 0.001 | 0 | 0.001 | 0.002 |
| 25 | BPI | 8.715 | 0.000 | 0.000 | 0.002 | 0.001 | 0 | 0.001 | 0.002 |
|  | API | 12.248 | 13.124 | 0.001 | 0.003 | 0.001 | 0 | 0.002 | 0.003 |
| 50 | BPI | 9.156 | 0.000 | 0.000 | 0.002 | 0.001 | 0 | 0.001 | 0.002 |
|  | API | 11.153 | 19.260 | 0.001 | 0.003 | 0.001 | 0 | 0.001 | 0.003 |
| 75 | BPI | 7.820 | 0.000 | 0.000 | 0.002 | 0.001 | 0 | 0.001 | 0.002 |
|  | API | 6.447 | 19.113 | 0.000 | 0.002 | 0.001 | 0 | 0.001 | 0.002 |
| 100 | BPI | 0 | 0 | 0 | 0 | 0 | 0 | 0 | 0 |
|  | API | 0 | 0 | 0 | 0 | 0 | 0 | 0 | 0 |

It can be seen from Table 10 that the Basic and the Advanced algorithms perform better than the Normal method for all the performance metrics. Since the non-edge priors are also considered, the Basic algorithm shows a reduction in the false positives when compared to the Normal method. However, the Advanced algorithm still predicts the least number of false positives. For example, at 10% edge priors, the Advanced algorithm predicts 12,017 wrong edges on an average, which is eight times lower than the number of false positives predicted by the Normal method, and 66 fewer false positives than the Basic algorithm. The Advanced algorithm, however, predicts 40 true positives lower than the Basic algorithm, due to which some performance metrics of the Basic algorithm is marginally better than the Advanced algorithm. As the percentage of the edge priors increase, the number of wrong edges identified by the Basic algorithm does not improve but the Advanced algorithm shows greater reduction in the number of wrong edges.

### 4.3 Performance of the different components of the algorithm

The Basic algorithm incorporates only the prior knowledge of interactions while the Advance algorithm, a) incorporates the prior knowledge of interactions, b) includes the prior knowledge of regulators, c) identifies and removes the wrong edges, and d) identifies the edge priors not supported by the data. In these four components, the identification of the edge priors not supported by data does not influence the results and is only for further studies in wet-lab. Moreover, in the networks used here, all the edge priors were supported by the data. Thus, only the first three components are analysed individually to identify their effectiveness in improving the performance of GRN inference.

#### 4.3.1 Prior knowledge of interactions

The prior knowledge of interactions consists of the edge priors and the non-edge priors in the *edgePrior* and the *nEdgeP rior* variables, respectively. The performance metrics of these variables are computed and compared with the Normal method to identify the improvements. As discussed in section 4.2, since the non-edge prior incorporation is studied only with the Net1, the result of *nEdgePrior* is available only for that network.

Tables 11, 12, and 13 show the performance metrics of the *edgePrior* in each of the three networks Net1, Net3, and Net4, respectively for the different percentage of edge priors. They all show similar results. As the edge prior% increases, the true positives, recall, F1-score, AUROC, and AUPR increase proportionately. The false positives for the *edgePrior* are always zero, as expected, because there are only known correct edges. Thus, the precision and specificity are 1. Table 14 shows the performance metrics of the *nEdgePrior* for the Net1. Since the *nEdgePrior* contains only the known non-edge priors, its performance metrics should have no true positives and the false positives should be equal to the number of non-edges known. The results in Table 14 are as expected. Further, the recall is also zero, which implies that no correct edges are identified, as required.

**Table 11:**
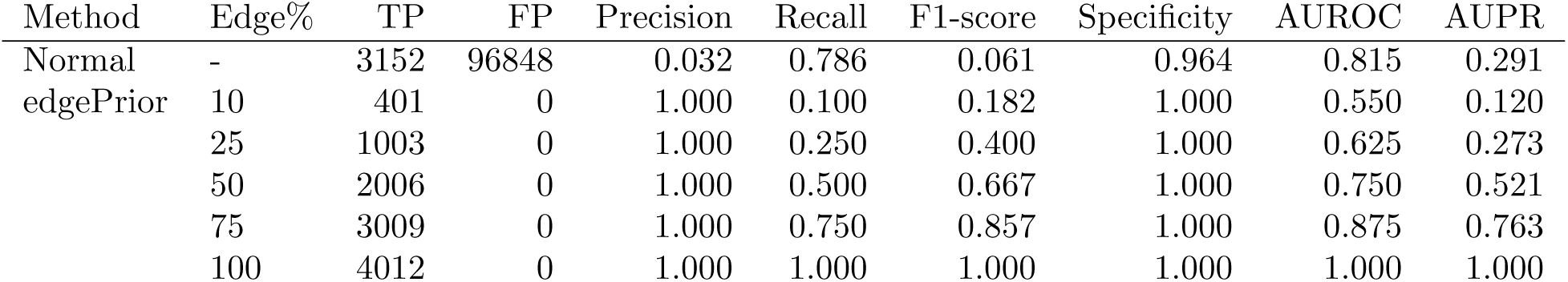
Performance metrics of Normal method and edge priors (*edgePrior* variable) for the Net1 network. In the performance metrics, TP is true positives and FP is false positives. The sample standard deviation was zero for all the conditions.

| Method | Edge% | TP | FP | Precision | Recall | F1-score | Specificity | AUROC | AUPR |
| --- | --- | --- | --- | --- | --- | --- | --- | --- | --- |
| Normal | - | 3152 | 96848 | 0.032 | 0.786 | 0.061 | 0.964 | 0.815 | 0.291 |
| edgePrior | 10 | 401 | 0 | 1.000 | 0.100 | 0.182 | 1.000 | 0.550 | 0.120 |
|  | 25 | 1003 | 0 | 1.000 | 0.250 | 0.400 | 1.000 | 0.625 | 0.273 |
|  | 50 | 2006 | 0 | 1.000 | 0.500 | 0.667 | 1.000 | 0.750 | 0.521 |
|  | 75 | 3009 | 0 | 1.000 | 0.750 | 0.857 | 1.000 | 0.875 | 0.763 |
|  | 100 | 4012 | 0 | 1.000 | 1.000 | 1.000 | 1.000 | 1.000 | 1.000 |

**Table 12:** Performance metrics of the non-edge priors (*nEdge Prior* variable) for the Net3 network. In the performance metrics, TP is true positives and FP is false positives. The sample standard deviation was zero for all the conditions.

| Method | Edge% | TP | FP | Precision | Recall | F1-score | Specificity | AUROC | AUPR |
| --- | --- | --- | --- | --- | --- | --- | --- | --- | --- |
| Normal | - | 614 | 99386 | 0.006 | 0.297 | 0.012 | 0.995 | 0.617 | 0.093 |
| edgePrior | 10 | 207 | 0 | 1 | 0.100 | 0.182 | 1.000 | 0.550 | 0.119 |
|  | 25 | 517 | 0 | 1 | 0.250 | 0.400 | 1.000 | 0.625 | 0.272 |
|  | 50 | 1033 | 0 | 1 | 0.500 | 0.667 | 1.000 | 0.750 | 0.520 |
|  | 75 | 1550 | 0 | 1 | 0.750 | 0.857 | 1.000 | 0.875 | 0.763 |
|  | 100 | 2066 | 0 | 1 | 1 | 1 | 1 | 1 | 1 |

**Table 13:** Performance metrics of Normal method and edge priors (*edgePrior* variable) for the Net4 network. In the performance metrics, TP is true positives and FP is false positives. The sample standard deviation was zero for all the conditions.

| Method | Edge% | TP | FP | Precision | Recall | F1-score | Specificity | AUROC | AUPR |
| --- | --- | --- | --- | --- | --- | --- | --- | --- | --- |
| Normal | - | 332 | 99668 | 0.003 | 0.084 | 0.006 | 0.997 | 0.518 | 0.021 |
| edgePrior | 10 | 394 | 0 | 1 | 0.100 | 0.182 | 1.000 | 0.550 | 0.124 |
|  | 25 | 985 | 0 | 1 | 0.250 | 0.400 | 1.000 | 0.625 | 0.277 |
|  | 50 | 1970 | 0 | 1 | 0.500 | 0.667 | 1.000 | 0.750 | 0.525 |
|  | 75 | 2955 | 0 | 1 | 0.750 | 0.857 | 1.000 | 0.875 | 0.765 |
|  | 100 | 3940 | 0 | 1 | 1 | 1 | 1 | 1 | 1 |

**Table 14:** Performance metrics of the non-edge priors (*nEdgeP rior* variable) for the Net1 network. In the performance metrics, TP is true positives and FP is false positives. The sample standard deviation was zero for all the conditions.

|  | Edge% | TP | FP | Recall |
| --- | --- | --- | --- | --- |
| nEdgePrior | 10 | 0 | 274380 | 0 |
|  | 25 | 0 | 274380 | 0 |
|  | 50 | 0 | 274380 | 0 |
|  | 75 | 0 | 274380 | 0 |
|  | 100 | 0 | 274380 | 0 |

Thus, incorporating the *edgePrior* can increase the true positives without increasing the wrong edges. This will also improve the other performance metrics. Incorporating the *nEdgePrior* will help to reduce the wrong edges without affecting the correct edges.

#### 4.3.2 Prior knowledge on regulators

The Advanced algorithm incorporates the knowledge of regulators by a) increasing the confidence weights of the inferred edges from the known transcription factors, and b) removing the inferred wrong edges from the non-transcription factors. Since non-transcription factors were not available, the second procedure was not performed. As discussed in section 2.7, the increased weights of the inferred edges from the known transcription factors are in the *ppEdgeTF*. By comparing the performance metrics of the *ppEdgeTF* to the Normal method the improvements given by the *ppEdgeTF* can be identified.

The performance metrics of the *ppEdgeTF* and the Normal method for Net1, Net3, and Net4 are in the tables 15, 16, and 17, respectively. It can be seen from the tables that the true positives detected for the *ppEdgeTF* are less than the Normal method. This is because the inferred edges from the known transcription factors are a subset of the total edges in the inferred network. Thus, the number of true positives in *ppEdgeTF* can never be larger than the true positives in the Normal method. The false positives are also much lower than the Normal method. Further, the precision, F1-score, and specificity are better than the Normal. This means that the procedure to increase the weights of transcription factor inferred edges, is identifying more correct edges with a very high true positive–to–false positive ratio compared to the Normal method. Thus, this component of the algorithm increases the weight of many true positives which will increase the performance of the Advanced algorithm.

**Table 15:**
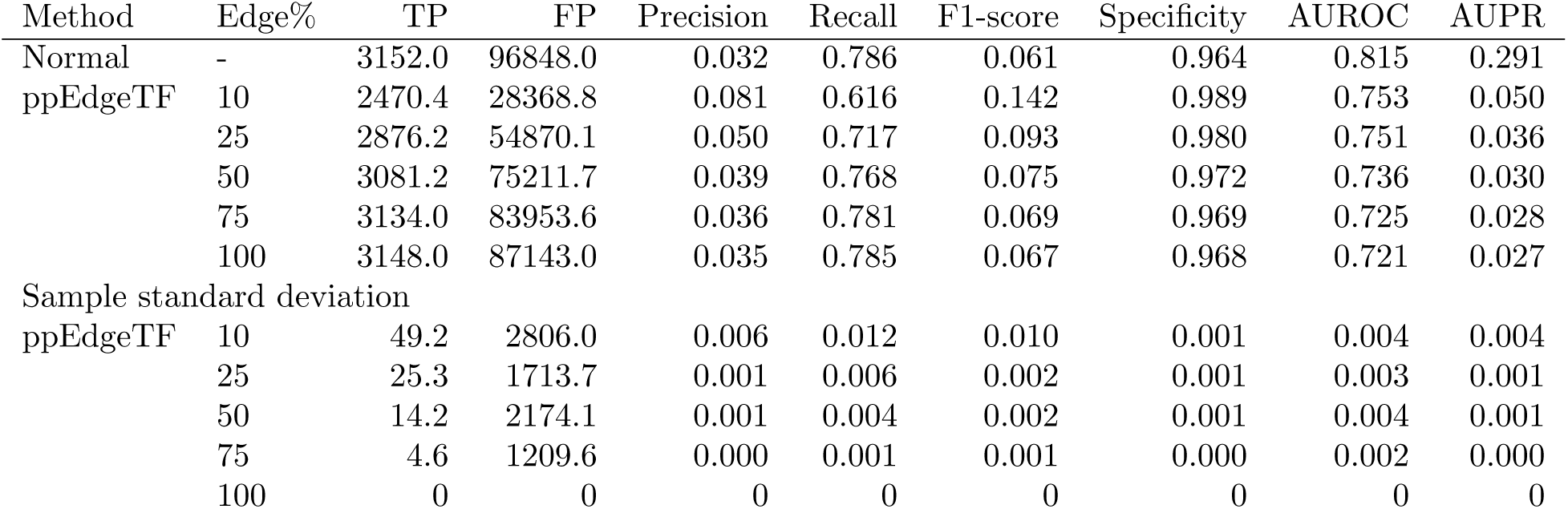
Performance metrics of Normal method and the *ppEdgeTF* for the Net1 network. In the performance metrics, TP is true positives and FP is false positives. A standard deviation equals to zero implies that its value is less than 0:001.

**Table 16:** Performance metrics of Normal method and the *ppEdgeTF* for the Net3 network. In the performance metrics, TP is true positives and FP is false positives. A standard deviation equals to zero implies that its value is less than 0.001.

| Method | Edge% | TP | FP | Precision | Recall | F1-score | Specificity | AUROC | AUPR |
| --- | --- | --- | --- | --- | --- | --- | --- | --- | --- |
| Normal | - | 614.0 | 99386.0 | 0.006 | 0.297 | 0.012 | 0.995 | 0.617 | 0.093 |
| ppEdgeTF | 10 | 356.8 | 13648.8 | 0.026 | 0.173 | 0.045 | 0.999 | 0.577 | 0.041 |
|  | 25 | 501.6 | 23561.1 | 0.021 | 0.243 | 0.039 | 0.999 | 0.604 | 0.040 |
|  | 50 | 565.9 | 32085.5 | 0.017 | 0.274 | 0.033 | 0.998 | 0.612 | 0.036 |
|  | 75 | 591.2 | 37352.9 | 0.016 | 0.286 | 0.030 | 0.998 | 0.614 | 0.034 |
|  | 100 | 604.0 | 41330.0 | 0.014 | 0.292 | 0.027 | 0.998 | 0.615 | 0.033 |
| Sample standard deviation |  |  |  |  |  |  |  |  |  |
| ppEdgeTF | 10 | 21.9 | 1504.3 | 0.002 | 0.011 | 0.003 | 0 | 0.004 | 0.002 |
|  | 25 | 13.7 | 1770.1 | 0.001 | 0.007 | 0.002 | 0 | 0.002 | 0.003 |
|  | 50 | 9.5 | 1491.9 | 0.001 | 0.005 | 0.001 | 0 | 0.002 | 0.001 |
|  | 75 | 2.5 | 1402.6 | 0.001 | 0.001 | 0.001 | 0 | 0.001 | 0.001 |
|  | 100 | 0 | 0 | 0 | 0 | 0 | 0 | 0 | 0 |

**Table 17:** Performance metrics of Normal method and the *ppEdgeTF* for the Net4 network. In the performance metrics, TP is true positives and FP is false positives. A standard deviation equals to zero implies that its value is less than 0:001.

| Method | Edge% | TP | FP | Precision | Recall | F1-score | Specificity | AUROC | AUPR |
| --- | --- | --- | --- | --- | --- | --- | --- | --- | --- |
| Normal | - | 332.0 | 99668.0 | 0.003 | 0.084 | 0.006 | 0.997 | 0.518 | 0.021 |
| ppEdgeTF | 10 | 287.0 | 16490.8 | 0.017 | 0.073 | 0.028 | 1.000 | 0.523 | 0.022 |
|  | 25 | 317.4 | 22562.9 | 0.014 | 0.081 | 0.024 | 0.999 | 0.523 | 0.021 |
|  | 50 | 327.3 | 26267.6 | 0.012 | 0.083 | 0.021 | 0.999 | 0.521 | 0.021 |
|  | 75 | 329.5 | 27626.0 | 0.012 | 0.084 | 0.021 | 0.999 | 0.520 | 0.021 |
|  | 100 | 331.0 | 28748.0 | 0.011 | 0.084 | 0.020 | 0.999 | 0.519 | 0.020 |
| Sample standard deviation |  |  |  |  |  |  |  |  |  |
| ppEdgeTF | 10 | 9.9 | 907.8 | 0.001 | 0.003 | 0.001 | 0 | 0.001 | 0 |
|  | 25 | 4.5 | 672.8 | 0 | 0.001 | 0.001 | 0 | 0.001 | 0 |
|  | 50 | 2.3 | 709.7 | 0 | 0.001 | 0.001 | 0 | 0.001 | 0 |
|  | 75 | 1.6 | 525.9 | 0 | 0 | 0 | 0 | 0 | 0 |
|  | 100 | 0 | 0 | 0 | 0 | 0 | 0 | 0 | 0 |

#### 4.3.3 Identifying and removing wrong edges

As discussed in section 2.7, the Advanced algorithm tries to identify and remove the wrong edges caused by the indirect interactions. The identified wrong edges are in the *ppNEdge*. When the prior knowledge of interactions are insufficient, the Advanced algorithm uses an approximate method to identify the wrong edges which was observed to cause a slight reduction in the number of true positives; while removing many false positives (see results in sections 4.2). Thus, computing the performance metrics of the *ppNEdge* will help to understand its effectiveness in identifying the wrong edges and improving the performance.

To be effective, the *ppNEdge* should: detect all or a large number of non-edges; detect no or minimum edges; and have a high non-edge–to–edge ratio. These conditions in the performance metric language implies that, for the *ppNEdge*, the number of true positives should be as low as possible, the number of false positives should be as high as possible, and the false positives–to–true positives ratio should be as high as possible.

Tables 18, 19, and 20 show the performance metrics of the *ppNEdge* in Net1, Net3, and Net4, respectively. It can be seen from the tables that all the three networks have similar results. The true positives are much lower than the false positives. The recall values are very low with their maximum values being 3.2%, 1.7%, and 1.0% for Net1, Net3, and Net4, respectively. This means that the number of true positives detected for each network is very low and is well below 4% of all the true positives. The number of false positives detected is large. The false positives–to–true positives ratio is more than 53, 57 and 94 for the Net1, Net3, and Net4, respectively for all the edge prior%. This means that, for the Net1, at least 53 false positives (wrong edges) are identified and removed for each true positive lost. Thus, identifying-and-removing-wrong-edge component is able to remove many wrong edges even when the prior knowledge is limited.

**Table 18:** The performance metrics of the *ppNEdge* that identifies and removes wrong edges for Net1 at different percentage of edge priors (Edge%). In the metrics TP is true positives and FP is false positives.

| Method | Edge% | TP | FP | Recall | FP/TP |
| --- | --- | --- | --- | --- | --- |
| ppNEdge | 10 | 25.6±6.6 | 1362.9±193.8 | 0.006±0.002 | 53.2 |
|  | 25 | 56.7±6.2 | 3369.6±235.5 | 0.014±0.002 | 59.4 |
|  | 50 | 92.8±5.2 | 6255.5±197.3 | 0.023±0.001 | 67.4 |
|  | 75 | 112.1±5.2 | 8547.7±212.7 | 0.028±0.001 | 76.3 |
|  | 100 | 127.0±0.0 | 10360.0±0.0 | 0.032±0.000 | 81.6 |

**Table 19:**
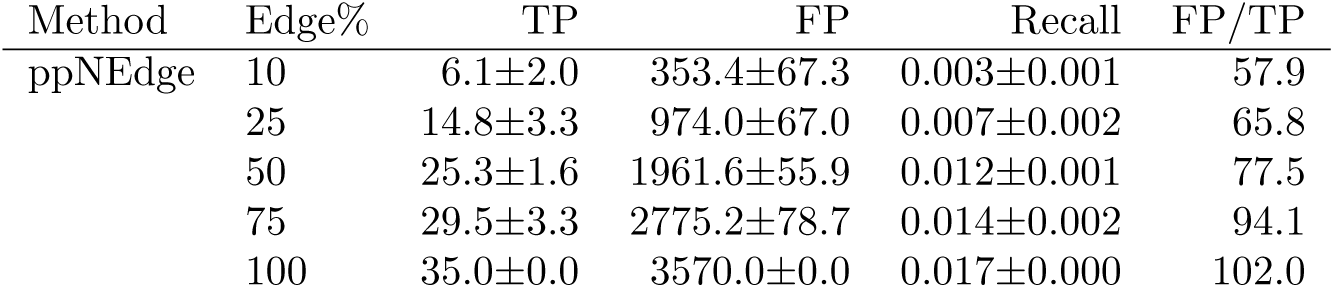
The performance metrics of the *ppNEdge* that identifies and removes wrong edges for Net3 at different percentage of edge priors (Edge%). In the metrics TP is true positives and FP is false positives.

**Table 20:** The performance metrics of the *ppNEdge* that identifies and removes wrong edges for Net4 at different percentages of edge priors (Edge%). In the metrics TP is true positives and FP is false positives.

| Method | Edge% | TP | FP | Recall | FP/TP |
| --- | --- | --- | --- | --- | --- |
| ppNEdge | 10 | 6.0±4.2 | 627.5±32.9 | 0.002±0.001 | 104.6 |
|  | 25 | 18.5±2.9 | 1746.0±77.6 | 0.005±0.001 | 94.4 |
|  | 50 | 27.8±1.8 | 3414.9±69.7 | 0.007±0.000 | 122.8 |
|  | 75 | 35.9±2.0 | 4856.9±55.4 | 0.009±0.000 | 135.3 |
|  | 100 | 40.0±0.0 | 6219.0±0.0 | 0.010±0.000 | 155.5 |

### 4.4 Discussion

The prior biological knowledge was incorporated into the inferred network obtained from a GRN inference method, that was the best-ranking method but was not known to be inherently capable of incorporating the prior knowledge. Currently, the prior knowledge can be incorporated mainly with the Bayesian network methods. However, these methods are not among the best-performing GRN inference methods (Marbach et al., 2012). Also, the current best-performing method cannot take advantage of the prior knowledge to improve its GRN inference accuracy. Since, the prior knowledge is incorporated in a post-inference step in this work, any GRN inference method can be used. This overcomes the limitation that only a few GRN inference methods can incorporate the prior knowledge.

Two post-inference prior algorithms were proposed, the Basic and the Advanced algorithms. The prior knowledge incorporation studies were performed on the widely used prokaryotic, eukaryotic, and in silico GRNs. The results of the studies showed that the proposed algorithms have much improved accuracy than the best-ranked GRN inference method from the DREAM5.

All the current GRN inference techniques based on the gene expression data infer edges due to the indirect interactions. This is because any edge that shows a statistical dependence at the gene expression level can be inferred. Moreover, it is not possible to differentiate statistical dependence and actual biological interaction due to the limitation in the data. The indirect interactions will be classified as wrong edges in the performance metrics. The Advanced algorithm can identify and remove the wrong edges due to the indirect interactions, when the prior knowledge of actual interactions are known. Thus, only the Advanced algorithm showed a reduction in the wrong edges when the non-edge priors were not available. Even when the non-edge priors were available, only the Advanced algorithm showed larger reduction in the wrong edges with the increasing quantity of the edge priors.

The four main components of a post-inference algorithm are a) incorporation of the prior knowledge of interactions, b) incorporation of the prior knowledge of regulators, c) identification and removal of the wrong edges, and d) shortlisting of the edge priors not supported by the data. These components have been separately analysed. The Basic algorithm uses only the first component while the Advanced algorithm uses all the four components. The prior knowledge of interactions consists of the edge and non-edge priors. Incorporating the edge priors increased the true positives without increasing the wrong edges, and incorporating the non-edge priors reduced the wrong edges without affecting the correct edges. Incorporating the prior knowledge of regulators consisted of increasing the weights of transcription factor inferred edges and removing the non-edges from the non-transcription factors. Increasing the weights of the known transcription factor inferred edges was found to increase the weights of many correct edges in the inferred network. Identifying and removing the wrong edge component was able to remove many wrong edges even when the prior knowledge was limited. The edge priors not supported by the data were detected using the MI metric and they could be interesting candidates for further study in wet-lab.

## 5 Conclusion

This study proposed that the prior biological knowledge can be incorporated using a post-inference step in a GRN inference method. Two post-inference prior algorithms — Basic and Advanced algorithms — have been developed and demonstrated. It is shown that the prior knowledge can be incorporated into any GRN inference technique with the post-inference prior algorithms. Both the algorithms show much better accuracy than the best-performing GRN inference method from the DREAM5. Further, it is also shown that by using the post-inference step, the wrong edges due to the indirect interactions can be identified and removed.

This work is significant because the prior knowledge integration is one of the common methods for overcoming the limitations of the high-throughput gene expression data based GRN inference (Chasman et al., 2016; Ghanbari et al., 2015; Gao and Wang, 2011; Hecker et al., 2009; Cooke et al., 2009; Hartemink et al., 2002). Further, as time progresses, the prior knowledge will also increase. Thus, inclusion of the prior knowledge along with the high-throughput data in the inference studies will be crucial.

The existing methods of incorporating the prior knowledge show better performance accuracy than the inference based on the data alone. Moreover, with a larger quantity of priors, their performance is further increased. The algorithms proposed here also show similar results. Moreover, one of the algorithms proposed here also reduces the wrong edges. Quantitatively comparing the existing methods of prior incorporation and the post-inference method of prior incorporation which is proposed here, is a subject for future study.

## Author contributions

AN and MC conceptualised the work. AN proposed and developed the algorithms, carried out the experiments, and analysed the results. AN and MC drafted the manuscript. All authors read and approved the final manuscript.

## Acknowledgements

AN performed part of this work in Pramod Wangikar’s lab. He thanks Pramod Wangikar for taking part in conceptualisation and some of the discussions on this work. He thanks Saranya Chandrasekharan, Kushal Satpute, Rose Davis, and Annesha Sengupta for general comments on the work and manuscript.

